# Aversive memories can be weakened during human sleep via the reactivation of positive interfering memories

**DOI:** 10.1101/2023.12.05.570072

**Authors:** Tao Xia, Danni Chen, Shengzi Zeng, Ziqing Yao, Jing Liu, Shaozheng Qin, Ken A. Paller, S. Gabriela Torres Platas, James W. Antony, Xiaoqing Hu

## Abstract

Recollecting painful or traumatic experiences can be deeply troubling. Sleep may offer an opportunity to reduce such suffering. We developed a procedure to weaken older aversive memories by reactivating newer positive memories during sleep. Participants viewed 48 nonsense words each paired with a unique aversive image, followed by an overnight sleep. In the next evening, participants learned associations between half of the words and additional positive images, creating interference. During the following non-rapid-eye-movement sleep, auditory memory cues were unobtrusively delivered. Upon waking, presenting cues associated with both aversive and positive images during sleep, as opposed to not presenting cues, weakened aversive memory recall while increasing positive memory intrusions. Substantiating these memory benefits, computational modeling revealed that cueing facilitated evidence accumulation toward positive affect judgments. Moreover, cue-elicited theta brain rhythms during sleep predominantly predicted the recall of positive memories. A noninvasive sleep intervention can thus modify aversive recollection and affective responses.

**Significance Statement:** Not all memories are welcome: recollection and involuntary intrusions of aversive memories can be debilitating. Prior research shows that reactivating positive memories benefits psychological health, and memory reactivation can be externally triggered during human sleep. Here, we introduced a novel sleep-based memory updating procedure to weaken older aversive memories by reactivating newer positive memories. We found that such reactivation not only weakened the recall of aversive memories, but also increased involuntary intrusion of positive memories. Moreover, reactivation enhanced positive affect judgments. Our findings open new avenues for weakening aversive or traumatic memories during human sleep.

## Introduction

Aversive memories can be overwhelming when they intrude on mnemonic awareness, impair cognitive function, and deteriorate mental health (Monfils & Holmes, 2018; Pitman et al., 2012). For many years, scientists have sought methods to help individuals manage these troubling memories (Anderson & Hulbert, 2021; Kredlow et al., 2016; Monfils & Holmes, 2018; Wixted, 2004). However, controlling aversive memories is challenging and effortful, due to their intense emotional charge and well-consolidated nature (Hu et al., 2017; Liu et al., 2016; Phelps & Hofmann, 2019; Vervliet et al., 2013). An alternative, less-studied route can be weakening aversive memories and even their affective responses during sleep, given that sleep influences both memory and emotion processing (Rasch & Born, 2013; Walker & van der Helm, 2009). Recent research has demonstrated that activating positive memories during wakefulness can reduce negative affect and depressive symptoms in humans, as well as alleviate depression-like behavior in rodents (Grella et al., 2022; Ramirez et al., 2015; Speer et al., 2021; Speer & Delgado, 2017; Wang et al., 2021). Here, we test a novel strategy to weaken aversive memories by introducing positive interfering memories, and then reactivating these memories during subsequent non-rapid-eye-movement (NREM) sleep.

Cross-species evidence suggests that memories of daily experiences are spontaneously reactivated during sleep, thus contributing to their consolidation (Klinzing et al., 2019; Nishida et al., 2009; Rasch & Born, 2013; Walker & van der Helm, 2009). Notably, memory consolidation can be selectively maneuvered by replaying memory-associated sensory cues during post-learning NREM sleep, a procedure known as targeted memory reactivation (TMR) (Cellini & Capuozzo, 2018; Hu et al., 2020; Oudiette & Paller, 2013; Paller et al., 2021). Although TMR is often used to strengthen memories, it also holds promises for weakening memories. When reactivating emotional memories during sleep in prior studies, results have been mixed. Some studies found that TMR strengthens emotional memories, others reported TMR benefits in subjective arousal reduction, while other studies found null effects on either memories or emotional responses (Ashton et al., 2018; Cairney et al., 2014; Hutchison et al., 2021; Lehmann et al., 2016; Pereira et al., 2022). Recently, research suggested that TMR could reactivate multiple memory traces during sleep, either strengthening or weakening episodic memories (Antony et al., 2018a; Joensen et al., 2022; Oyarzún et al., 2017). To the best of our knowledge, no studies have been designed to weaken previously acquired, older aversive memories during sleep by reactivating their corresponding positive memories.

Examining cue-elicited neural activity during TMR could illuminate neural mechanisms that drive memory change, potentially deepening our understanding of reactivation of positive interfering memory during sleep. Specifically, slow-wave sleep (SWS) and spindle-related sigma power have consistently been shown to play crucial roles in TMR-induced memory benefits (Antony, et al., 2018b; Cairney et al., 2018; Creery et al., 2015; Liu et al., 2023; Schreiner et al., 2018; Xia, et al., 2023a). In the case of reactivating interfering memories, enhanced beta (16-30 Hz) activity may implicate memory interference during sleep and predict post-sleep forgetting (Antony et al., 2018a; Oyarzún et al., 2017). Moreover, 4-8 Hz theta activity has been associated with emotional processing and emotional memory reactivation during sleep (Canales-Johnson et al., 2020; Lehmann et al., 2016; Xia et al., 2023b). Here, we focused on theta and beta activity to elucidate the neural mechanisms underlying the reactivation of interfering positive memories during sleep.

We hypothesized that older aversive memories could be weakened by reactivating their corresponding positive memories during NREM sleep. To test this hypothesis, we designed a multi-day procedure (Figure 1). Participants encoded aversive memories on Day 1 followed by an overnight sleep for memories to consolidate. On Day 2 evening, participants encoded positive memories that shared the same cues as half of the older aversive memories, thereby producing interference. During subsequent NREM sleep, we replayed memory cuesassociated with both aversive and positive images to weaken the older aversive memories and affective responses. Recognizing multiple expressions of emotional memory (Phelps & Hofmann, 2019), we assessed both memory (voluntary recall and involuntary intrusions) and affective responses (subjective ratings and speeded affect judgments) related to both aversive and positive memories. Including these measures allowed a systematic examination of TMR impact on various expressions of aversive memories.

**Figure 1.**
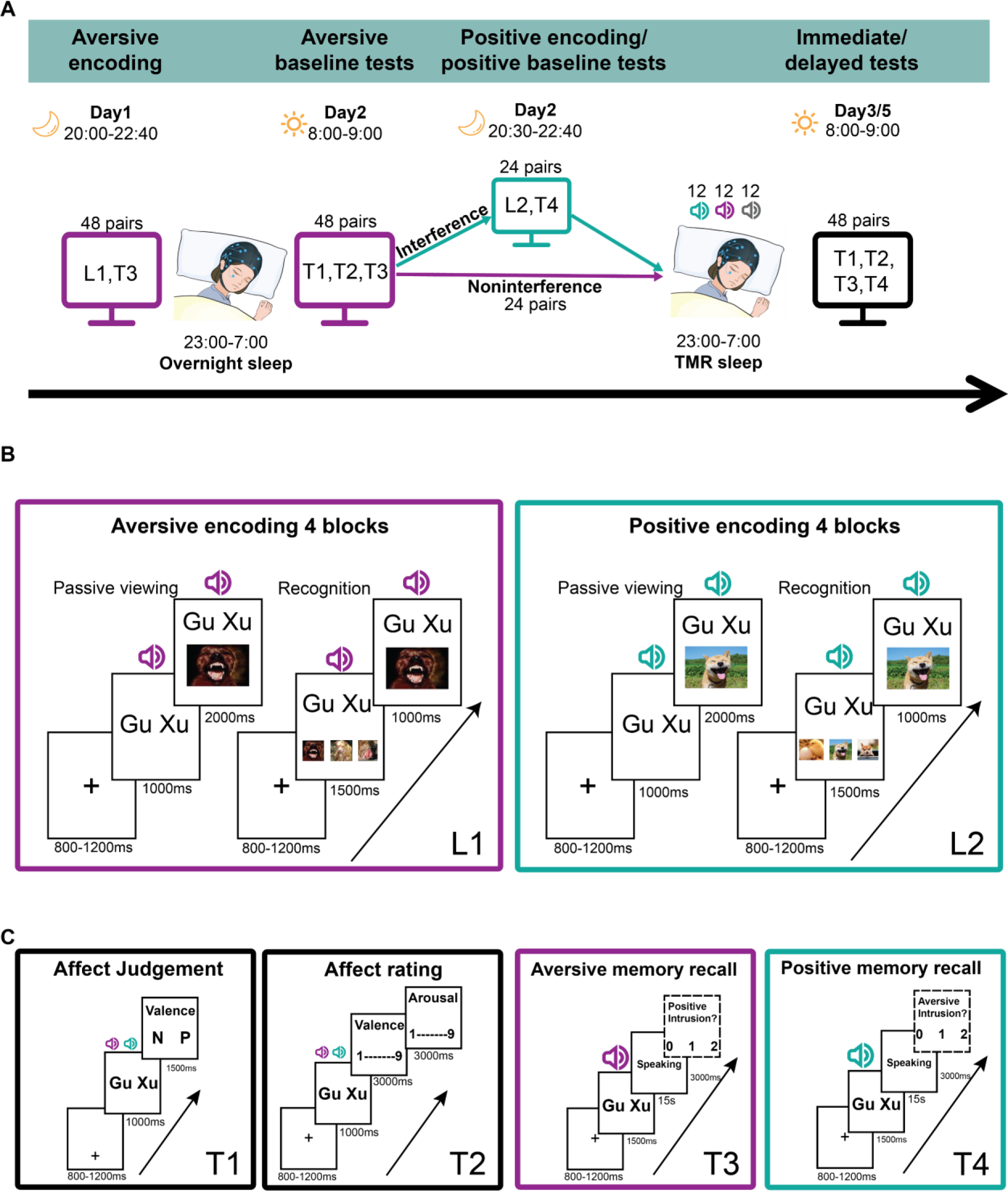
Experimental procedure. (A) On Day 1, participants learned 48 unique pseudoword-image pairs with only aversive images. On Day 2, 24 of the same pseudowords were paired with positive images, introducing interfering memories. L1 and L2 denote the learning task with aversive or positive images, respectively. The 48 images were comprised of 12 images from four content categories (animal, baby, people, and scene), with two of the categories (i.e., 24 word-image pairs) selected randomly for L2. Two conditions were thus created, the interference condition (pseudowords associated with both an aversive and a positive image) and the noniterference condition (pseudowords associated with only an aversive image). During a TMR sleep session, 12 memory cues from the interference condition (green sound icon) and 12 from the noninterference condition (purple sound icon), were presented to sleeping participants during NREM sleep. Additionally, 12 novel pseudowords (grey sound icon) that were not paired with any images during the experiment were played as control sounds. TMR cues were presented in a randomized block manner. T1, T2, T3, and T4 denote testing tasks as schematized in panel C. (B) The left panel (purple box) represents aversive encoding on Day 1 and the right panel (green box) positive encoding on Day 2. Each encoding round included passive viewing followed by recognition-with-feedback tests, wherein three previously viewed images were presented as options. Participants completed four encoding blocks to achieve high encoding accuracy. (C) The left two panels (black boxes) represent affect-judgment and affect-rating tasks, and the right two panels (purple and green boxes) illustrate aversive and positive cued-recall tasks. During each cued recall task, participants also reported the presence or absence of intrusions. Importantly, no positive intrusions were assessed during aversive recall on Day 1 and the morning of Day 2, as positive memories had not yet been acquired at these time points.

Our experimental design, with positive memories acquired just prior to TMR sleep, may facilitate their reactivation during TMR due to temporal proximity (Oyarzún et al., 2017; Seibold et al., 2018). Moreover, this design offers an intriguing context for exploring how to weaken older aversive memories. We found that memory cueing during NREM sleep impaired subsequent recall of aversive memories, together with increased involuntary intrusions of positive memories. Regarding affect changes, cueing increased positive affect judgments toward the cues, and facilitated evidence accumulation toward positive judgments as revealed by the drift diffusion model. Examining cue-elicited EEG activity suggested that TMR preferentially reactivated positive interfering memories during NREM sleep, as evidenced by cue-elicited theta power predicted subsequent positive memories.

## Results

### Effective learning of aversive associations

Given that we aimed to weaken the earlier acquired aversive memories, which may have been partially consolidated during an overnight night of sleep, we first examined the recognition performance from Day 1 evening. Over four blocks of viewing and recognition-with-feedback tests, accuracy increased (Figure 2A) while reaction times (RTs) decreased (Figure 2B). All participants achieved at least 75% correct recognition accuracy in the final block.

**Figure 2.**
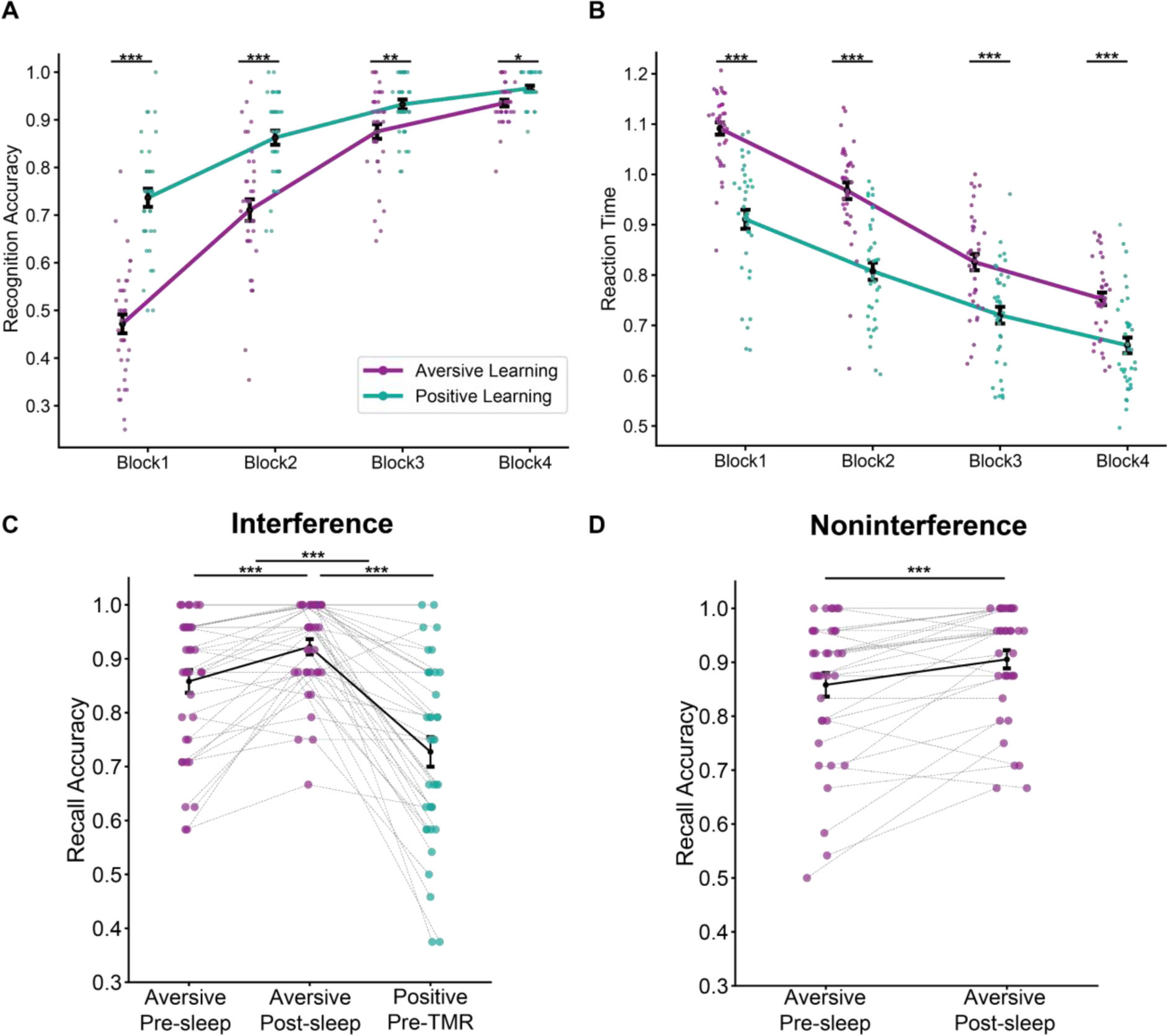
Comparing aversive and positive learning on Day 1 and Day 2. Results of aversive and positive learning on Day 1 and Day 2, respectively. Participants demonstrated significantly higher recognition accuracy (A) and faster response times (B) for positive recognition on Day 2 compared to aversive recognition on Day 1. (C) Aversive recall accuracy for pre- and post-sleep tests (24 items), and positive memory accuracy before the TMR sleep session in the interference condition (24 items). (D) Aversive recall accuracy during pre-overnight and post-overnight sleep tests in the noninterference condition (24 items). *** p < 0.001, ** p < 0.01, * p < 0.05, error bars represent standard errors.

Next, we examined recall of these aversive memories after learning and following sleep, by comparing pre- and post-sleep cued recall of aversive memories. Recall required participants to verbally describe the image after seeing the corresponding pseudoword, and then these descriptions were scored by raters to determine whether the image was correctly recalled for each trial (see Methods for details). A comparison of pre-versus post-sleep cued recall showed that accuracy was significantly superior after sleep, both in the interference and noninterference conditions (Interference: pre-sleep 0.86 ± 0.02; post-sleep 0.92 ± 0.01; paired-sample t-test, t(36) = 4.46, p < 0.001, **d** = 0.73; Noninterference: pre-sleep 0.86 ± 0.02; post-sleep 0.91 ± 0.02; t(36) = 3.78, p < 0.001, **d** = 0.62, Figure 2CD). In subsequent analyses, we used Day 2 morning cued recall accuracy as the aversive memory baseline performance.

### Effective learning of positive associations, producing interference

On Day 2 evening, participants learned associations between positive images and half of the pseudowords from Day 1, producing interference. Across learning blocks, participants’ recognition accuracy for positive memories was significantly higher than for aversive memories across all four blocks (positive vs. aversive recognition: 0.74 - 0.97 vs. 0.47 - 0.94 across blocks 1 to 4, t(36)s >3.44, p_corrected_s = 0.029, Figure 2A). Participants were also faster in correct recognitions of positive than aversive memories (positive vs. aversive: 0.91 - 0.66s vs. 1.09 – 0.75 across blocks 1 to 4, t(36)s > 6.52, p_corrected_s < 0.001, Figure 2B). High recognitions indicated that participants successfully learned the positive associations, enabling TMR to reactivate these memories and weaken corresponding aversive memories.

However, participants’ recall accuracy of positive memories on Day 2 evening (Mean ± SE: 0.73 ± 0.03) was significantly lower when compared to the recall of corresponding aversive memories on Day 1 evening (Mean ± SE: 0.86 ± 0.02; t(36) = 5.00, p_corrected_ < 0.001), and on Day 2 morning (Mean ± SE: 0.92 ± 0.01; t(36) = 7.37, p_corrected_ < 0.001; Figure 2 C). Notably, among the positive forgotten items (∼27% across participants), 87% of them were remembered during the aversive memory baseline test, while 90% of them were correctly recognized during the last block of positive encoding.

In summary, we found that both aversive and positive memories were well recognized during learning trials. Moreover, aversive memories were likely consolidated during the first overnight sleep. For positive memories, participants showed higher recognition accuracy during positive encoding blocks; but lower verbal recall of positive memories compared to the aversive memories. These results suggested that the earlier acquired aversive memories specifically interfered with the retrieval, rather than recognition, of positive interfering memories (i.e., output interference, Dyne et al., 1990; Runquist, 1975; Verde, 2004).

### TMR weakened older aversive memories and selectively improved positive memories

To answer our primary research question about weakening aversive memories, we examined TMR effects on the recall of earlier acquired aversive memories (i.e., T3 on Day 2, Day 3, and 5; for descriptives, see Table S1). We analyzed item-level aversive memory accuracy (correctly recalled or not) using a Bayesian linear mixed model (BLMM), given that this method allowed examining individual memory items and is well-suited for handling hierarchical data structures with large numbers of random effects and varying slopes (Sorensen et al., 2016). In this model, TMR (Cued vs. Uncued) and interference (Interference vs. Noninterference) were treated as fixed effects, while baseline aversive memory accuracy (Remembered vs. Forgotten) and time (Immediate vs. Delayed) were included as covariates. We incorporated time as a covariate rather than a fixed factor, as model comparisons indicated that the current model was superior to alternative models (i.e., treating time as a fixed factor, see Table S2 for model comparison). For consistency, all main results were analyzed with time treated as a covariate. Additionally, we provided the results that treated time as a fixed factor, revealing that TMR effects were more evident observed during the delayed test (see Table S3 for immediate and delayed results).

Results showed that in the interference condition, cueing (cued vs. uncued) reduced the recall of aversive memories (median*_diff_* = −0.44, 95% HDI [−0.82, −0.11], Figure 3A). However, in the noninterference condition, the cueing effect was not significant (median*_diff_* = 0.05, 95% HDI [−0.53, 0.64], Figure 3B). We next examined the TMR effects on positive interfering memories, using TMR as fixed factor and positive memory baseline accuracy and time as covariates. The results revealed a non-significant TMR effect on the recall of positive interfering memories (median*_diff_* = −0.18, 95% HDI [−0.23, 0.65]).

**Figure 3.**
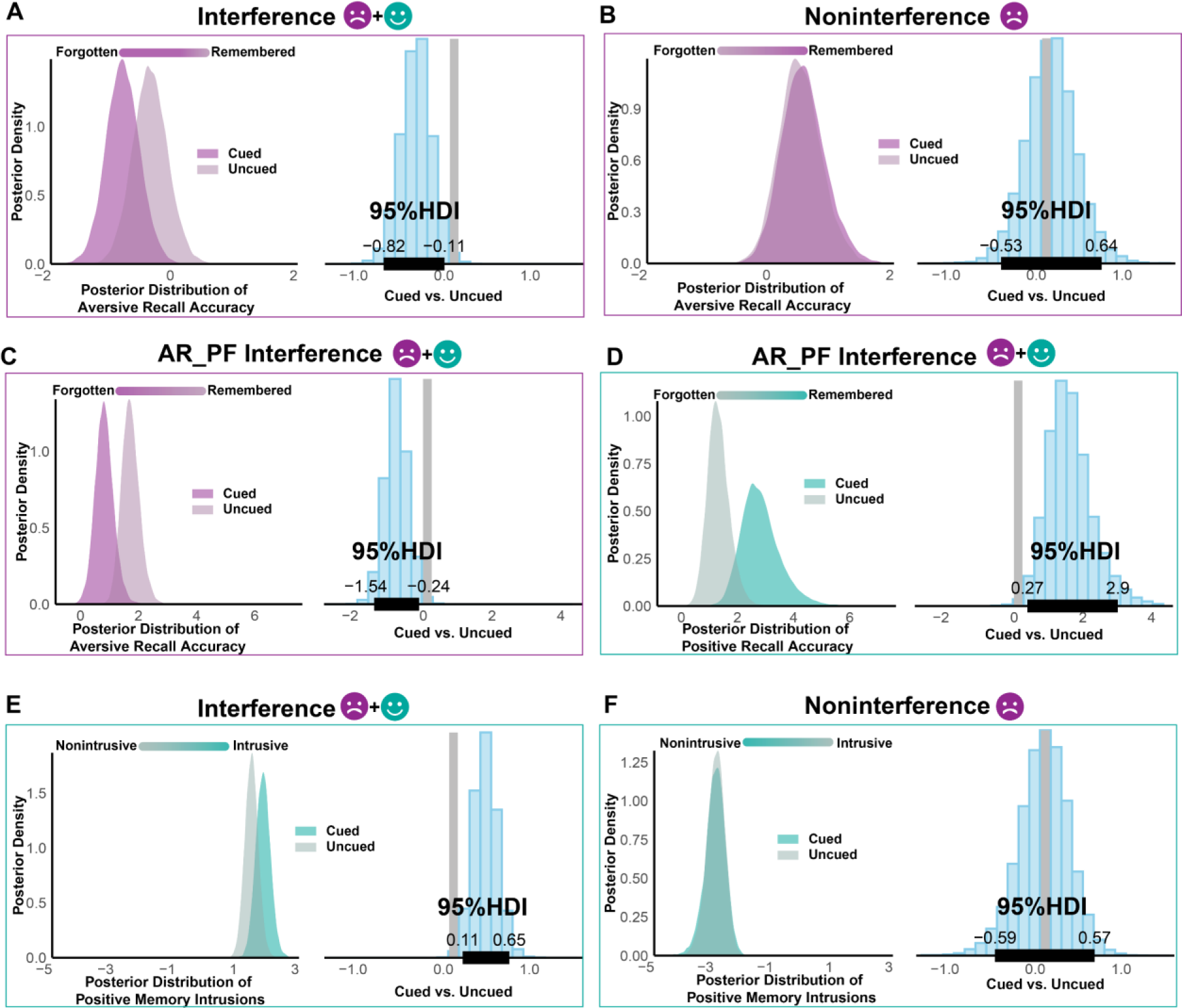
TMR weakens earlier acquired aversive memories and strengthens positive interfering memories in the interference condition. Item-level aversive memory, positive memory, and positive memory intrusions were analyzed using a Bayesian linear mixed model (BLMM). (A) TMR cueing in the interference condition reduced recall accuracy of aversive memories, (B) but not in the noninterference condition. (C-D) When we focused on aversive remembered yet positive forgotten (AR_PF) items before the TMR session, TMR not only weakened aversive memories (C) but also strengthened positive memories (D). (E-F) TMR induced more positive memory intrusions in the interference (E) condition but not in the noninterference (F) condition. In all panels, purple boxes indicate aversive dependent measures, green boxes represent positive dependent measures. Within each panel, the left-side density plots are derived from actual data and display the fitted data of the posterior distribution for the respective dependent variables (ABC: aversive memory recall accuracy; D: positive memory recall accuracy; EF: positive memory intrusions). The right-side histogram plots illustrate the contrast between cued and uncued conditions, with horizontal black lines representing the 95% highest density interval (HDI), and vertical gray lines denoting 0. If 0 does not overlap with the 95% HDI, the result is considered significant.

The above results suggested that TMR weakened aversive memories while not influencing positive interfering memories in the interference condition. We further examined the concurrent influence of TMR on both aversive and positive memories in the interference condition in a single model, with TMR (Cued vs. Uncued) and valence (Aversive vs. Positive memories) as fixed factors and time and baseline accuracy as covariates. Collaborating with previously mentioned results, we found that cueing significantly decreased aversive memory accuracy (Cued vs. Uncued, median*_diff_* = −0.34, 95% HDI [−0.68, −0.01]) without influencing positive memory accuracy (Cued vs. Uncued, median*_diff_* = 0.16, 95% HDI [−0.28, 0.62]).

Given that older aversive memories interfered with recall of corresponding positive memories during wakefulness, we hypothesized that these positive memories might be more susceptible to TMR. To investigate this possibility, we identified memory items that were forgotten during the Day 2 evening positive recall test but remembered during the Day 2 morning aversive memory recall test (Aversive remembered_Positive forgotten or AR_PF, ∼23.6% of all items across participants). We incorporated these items into the BLMM, wherein TMR (Cued vs. Uncued) and valence (Aversive vs. Positive memories) served as fixed factors, and time functioned as a covariate.

Results showed that for these memory items, cueing significantly reduced aversive memory accuracies (cued vs. uncued, median*_diff_* = −0.91, 95% HDI [−1.54, −0.24], Figure 3C) while increased positive memory accuracies (cued vs. uncued, median*_diff_* = 1.44, 95% HDI [0.27, 2.90], Figure 3D). This effect was specific to these aversive remembered_positive forgotten items, because the same analyses on other categories (aversive remembered_positive remembered, aversive forgotten_positive forgotten, aversive forgotten/positive remembered) did not show any cueing effect on aversive (0.29 < median*_diff_* < 3.36, all HDIs overlapped with 0) or positive memories (0.20 < median*_diff_* < 2.64, all HDIs overlapped with 0). These findings suggested that TMR weakened aversive memories while improving positive interfering memories, yet only when aversive memories had interfered the recall of the positive memories.

### TMR increased intrusions of positive memory during aversive recall

Besides voluntary recall, we employed the BLMM to examine the TMR effects on positive memory intrusions during aversive memory recall (T3), while treating time as a covariate for consistency (see Table S2 for immediate and delayed results separately). Note that for positive memory intrusion, there was no baseline performance because participants had not yet learned the positive memories on Day 2 morning. Therefore, positive intrusion was only examined on Day 3 and Day 5. The results revealed that cueing induced more positive memory intrusions in the interference condition (median*_diff_* = 0.37, 95% HDI [0.11, 0.65], Figure 3E) but not in the noninterference condition (median*_diff_* = −0.01, 95% HDI [−0.59, 0.57], Figure 3F). In contrast, cueing did not impact aversive memory intrusions during positive memory recall (T4) on Day 3 and Day 5 (median*_diff_* = −0.03, 95% HDI [−0.40, 0.33]). Together, we found that TMR enhanced positive intrusions during aversive memory recall, but did not influence aversive intrusions during positive memory recall.

### TMR increased positive affect judgments: behavioral and computational evidence

In addition to memory changes, we were interested in TMR’s impact on affect changes. For the speeded affect judgment task, we calculated the positive judgment ratios for each item (i.e., the total number of positive judgments divided by the total numbers of judgments for the specific item) at three time points (i.e., Day 2 morning, Day 3 morning, and Day 5 morning). We next calculated the positivity change scores (Day 3/5 morning minus Day 2 morning), and submitted it to BLMM to examine how TMR and interference impacted the positivity change scores using time as a covariate (see Methods for model details). Our findings revealed that compared to uncued items, cueing increased positivity change scores in the interference (median*_diff_* = 0.05, 95% HDI [0.01, 0.10], Figure 4A) but not in the noninterference condition (median*_diff_* = 0.01, 95%HDI [−0.04, 0.06], Figure 4B). The same analysis on RT changes did not yield significant differences in either interference (median*_diff_* = 0.02, 95%HDI [−0.004, 0.04]) or noninterference condition (median*_diff_* = 0.02, 95%HDI [−0.005, 0.04]). For the affect rating task, the same analysis was conducted on valence and arousal changes, which did not yield any significant results between cued and uncued conditions (−0.070 < median*_diff_* < 0.093, all HDIs overlapped with 0).

**Figure 4.**
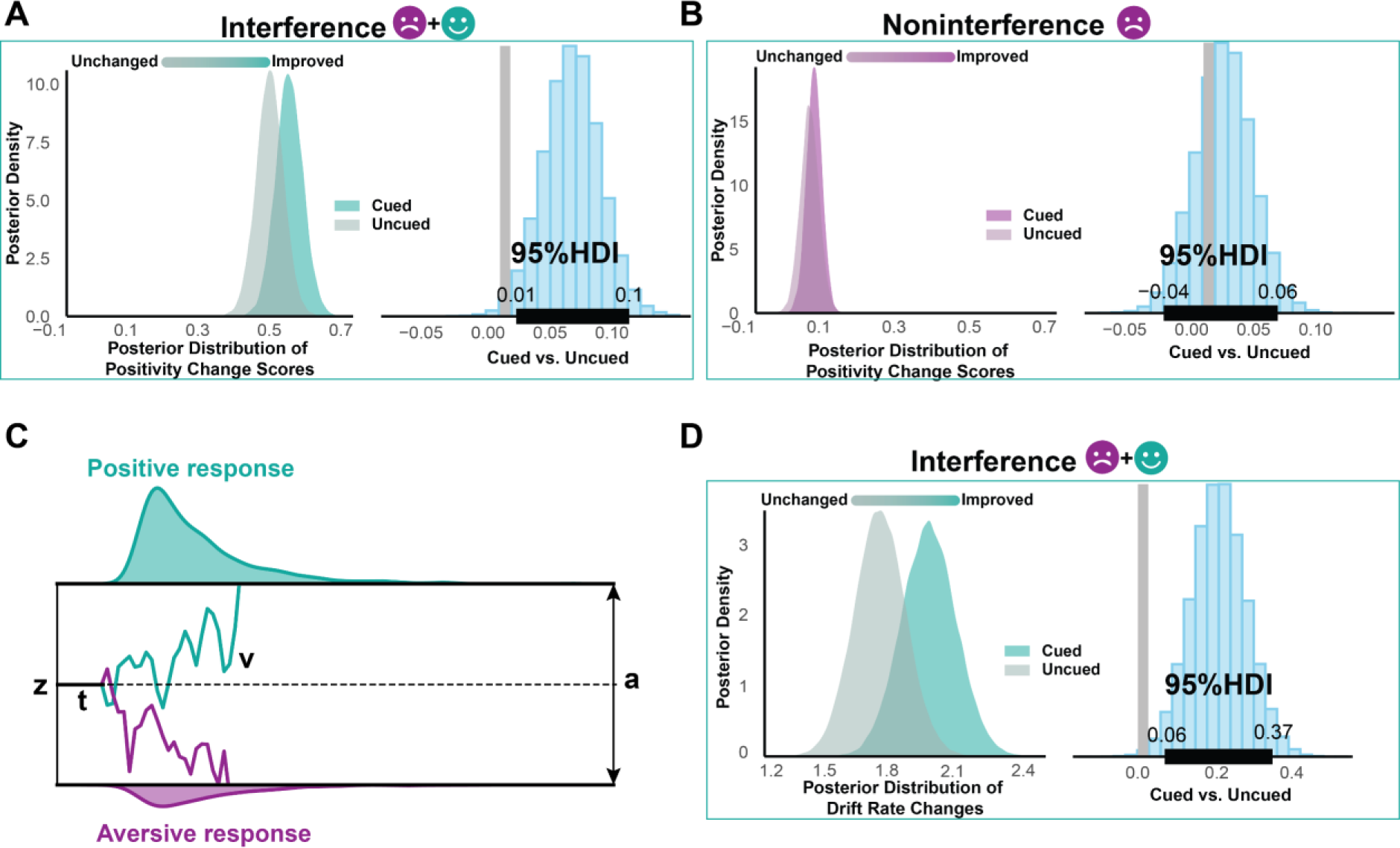
Computational modeling of affect judgments. Item-level positivity change scores were assessed using a BLMM. (AB) TMR increased positivity changes in the interference condition (A) but not in the noninterference condition (B). (C) Illustrates the general processing of the hierarchical drift diffusion model (HDDM). The main parameters of the model include the starting point z, the drift rate v, the decision boundary a, and the non-decision time t, which determine when sampled sensory evidence (green and purple) leads to a choice that is positive (green, upper boundary) or aversive (purple, lower boundary). (D) TMR facilitated evidence accumulation toward positive response in affect judgment task. The drift rate was calculated using the HDDM regression module. Within panels AB and D, the left-side density plots display the posterior distribution of the corresponding dependent variables (e.g. positivity change scores and drift rate change) in cued and uncued conditions. Figure AB is based on the actual data depicting positivity change scores, whereas Figure D is derived from the HDDM model representing drift rate changes, with the displayed distribution illustrating the fitted data of posterior distributions.The right-side histogram plots represent the contrast between cued and uncued conditions, with horizontal black lines indicating the 95% highest density interval (HDI), and vertical gray lines denoting 0. If 0 does not fall within the 95% HDI, the result is deemed significant.

To further delineate how TMR may impact the cognitive processing underlying the binary speeded affect judgments, we used a drift diffusion model (DDM, Ratcliff et al., 2004, 2016; White et al., 2009). The DDM is a well-established computational method examining optimal processes in a binary task within a sequential sampling framework. The decision-making process in the DDM consists of four key parameters: the starting point (*z*) between the two choice boundaries, indicating a pre-decision bias; the non-decision time (*t*), accounting for factors unrelated to decision-making; the noisy drift process (*v*), representing information accumulation toward the preferred choice boundary; and the choice boundaries (*a*), which signify the completion of the decision-making process when the accumulated evidence reaches either one of them (Figure 4C).

We hypothesized that TMR would influence the starting point (*z*) and drift rate (*v*) in the interference condition. To test this, we estimated *v*, *z*, and *vz* separately across different conditions in three different models, while estimating *a* and *t* at the participant level. We implemented a hierarchical Bayesian approach through the hierarchical drift diffusion model (HDDM), which offers more robustness and tolerance for low trial numbers (Wiecki et al., 2013). In our analysis, we incorporated TMR, time, and their interaction into the regression model of the HDDM (see methods for model details). Model comparison results suggested that the *v* model ranked highest. We thus focus our results on the drift rate.

Regarding the drift rate *v*, on Day 2 morning aversive baseline test, participants showed faster evidence accumulation speed for aversive than positive responses (median*_diff_* = −1.55, 95% HDI [−1.84, −1.28], indicating that participants effectively learned the aversive associations. On Day 3 morning following the TMR, participants showed faster evidence accumulation speed for positive than for aversive responses (median*_diff_* = 0.60, 95% HDI [0.21, 0.98]), though this effect was not observed during the Day 5 delayed test (median*_diff_* = −0.19, 95% HDI [−0.62, 0.23]).

Critical to our research question, we were particularly interested in how cueing changed drift rate from baseline to immediate and delayed tests. In line with previous analyses, we averaged the results of immediate and delayed tests and calculated the drift rate change from pre-sleep to post-sleep (see Figure S1 for immediate and delayed results separately). Our analysis revealed that participants were faster in accumulating evidence toward making positive judgments for cued items than uncued items from baseline to immediate and delayed tests (cued vs. uncued, median*_diff_* = 0.22, 95% HDI [0.06, 0.37], Figure 4D).

### Unraveling cue-elicited EEG responses during sleep

Our behavioral findings indicated that TMR weakened earlier acquired aversive memories while increasing positive memory intrusions in the interference condition. To examine how TMR reactivated aversive and positive memories during NREM sleep, we extracted cue-locked, time-frequency resolved EEG responses in the interference and noninterference conditions, and compared them with the EEG responses elicited by control sounds.

We found that when compared to the control sounds that did not involve any memory pairs before sleep, both interference and noninterference memory cues increased EEG power across the delta, theta, sigma, and beta bands in frontal and central areas (p_cluster_s < 0.01, corrected for multiple comparisons across time, frequency, and space, Figure 5AB, CD). However, when contrasting interference with noninterference memory cues, we did not identify any significant clusters (p_cluster_s > 0.05). These findings suggested that delta-theta and sigma-beta power increases may indicate memory reactivation during sleep.

We next examined whether cue-elicited theta and beta power were associated with subsequent memory accuracies (i.e., remembered vs. forgotten) for individual positive or aversive stimulus in the interference condition, given the relationship between theta activity and emotional processing (Nishida et al., 2009), and between beta activity and memory interference during sleep (Antony et al., 2018; Oyarzún et al., 2017). Employing BLMM across all channels revealed that the cue-elicited theta power over the right central-parietal region (FC5, C2, C4, CP2, CP4, TP7) was significantly higher for subsequently remembered positive memories than for forgotten positive memories (median*_diff_* = 1.39, 95% HDI [0.32, 2.43], Figure 5E). For aversive memories, a few channels’ (Fp2, F6, C5) theta power was higher for remembered than forgotten aversive memories (median*_diff_* = 1.04, 95% HDI [0.16, 1.86],; Figure 5F). In contrast, we found no relationships between cue-elicited beta power and positive or aversive memories were found (median*_diff_*s < 0.17, all HDIs overlapped with 0; see Figure 5G, H). In addition to investigating the subsequent memory effect on theta power increase, we also examined the subsequent effect of theta and sigma power decrease; however, no significant results were found (theta: 0.001 < median*_diff_*s < 0.099; Sigma: −0.008 < median*_diff_*s < 0.015; all HDIs overlapped with 0, see Figure S4).

**Figure 5.**
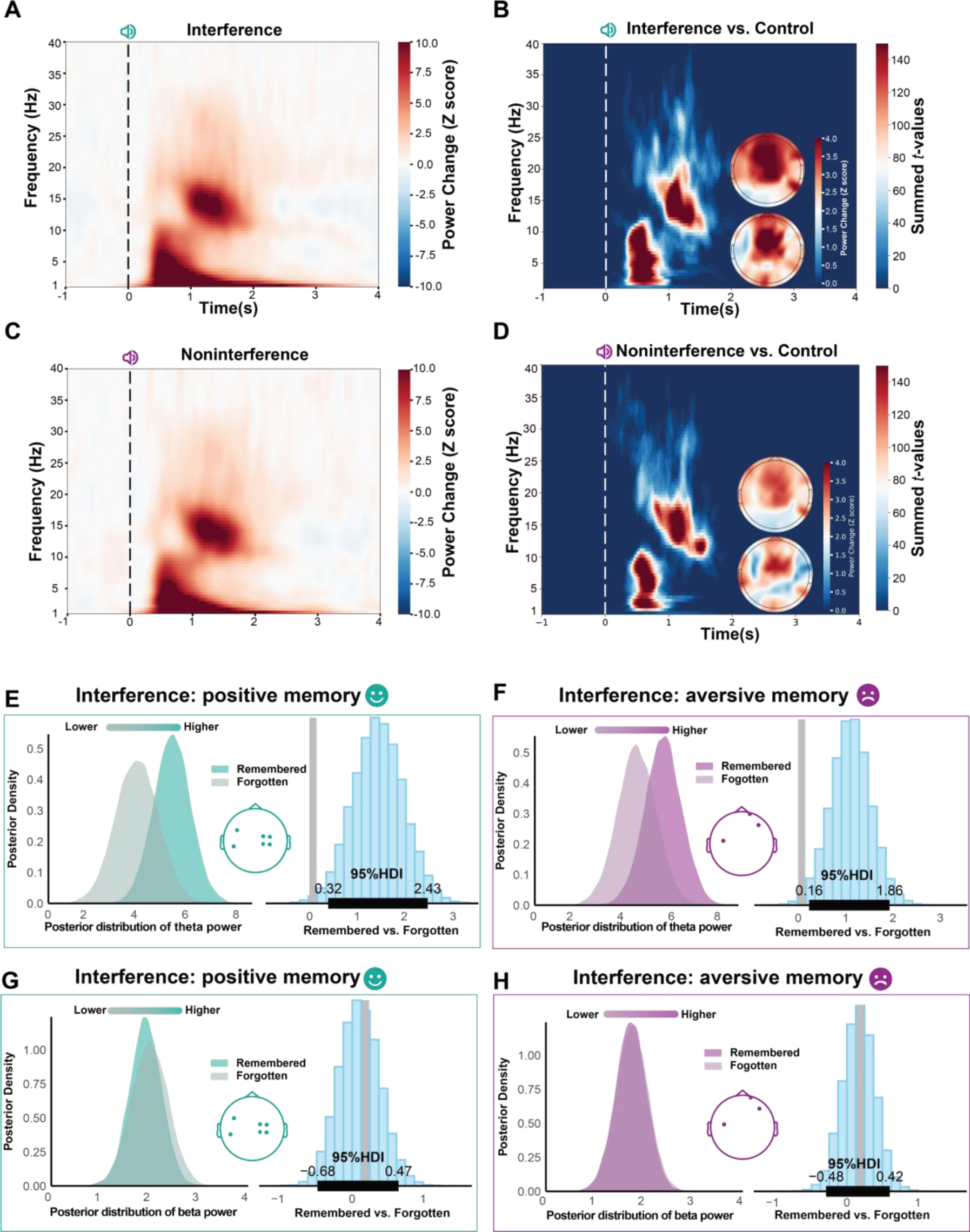
EEG response toward interference and noninterference cues. Time-frequency results were averaged across all trials and participants over all electrodes in the (A) interference and (C) noninterference condition. The power differences between memory cues (B: interference, D: noninterference) and control sounds are illustrated in the image plot, which displays the t-map from a cluster-based permutation test across time points, frequency bands, and channels. The bottom topographical plot shows the raw values (Z-scored) of the delta-theta band, while the upper plot presents the raw values of the sigma-beta bands. (E, F) Cue-elicited theta power for subsequently remembered positive and aversive items was significantly greater than that of forgotten items across different channels; (G, H) while no beta power differences were found between remembered and forgotten positive or aversive memories. Theta power results are averaged across significant channels for positive and aversive memory items. For consistency, beta power results are averaged using the same channels as from the theta results. In all panels, green outlines represent positive memory outcomes; and purple outlines represent aversive memory outcomes. Within panels EFGH, the left-side density plots are derived from actual data and display the fitted data of the posterior distribution of theta and beta power activity in remembered and forgotten conditions, with the right-side histogram plots illustrating the contrast between remembered and forgotten conditions. Horizontal black lines represent the 95% highest density interval (HDI), and vertical gray lines denote 0. If 0 does not overlap with the 95% HDI, the result is considered as significant.

### Cue-elicited theta power predicted memory changes

Our previous results revealed that cue-elicited theta power was associated with subsequent memory accuracies, suggesting that cue-elicited theta power may track cue-triggered memory reactivation during sleep. In light of these findings, we proceeded to investigate the relationship between cue-elicited theta power and TMR effects on both aversive and positive memories at post-sleep tests. To quantify the relationship between EEG power increases and item-level aversive and positive memories, we employed BLMM with EEG power and valence (positive vs. aversive memories) as fixed factors, incorporating baseline memory accuracy and time as covariates. Single-item EEG activity was extracted from significant clusters spanning across all channels and was subsequently averaged across time points and frequency bands within these clusters. This analysis incorporated the single-item EEG power from each frequency band into the model, and was repeated for each channel, allowing for a thorough examination of possible EEG-behavior relationships.

Results revealed that cue-elicited theta power positively predicted both positive (median*_diff_*s > 0.05, all HDIs did not overlap with 0, see Figure 6A) and aversive (median*_diff_*s > 0.03, see Figure 6 B) memory recall, whereas cue-elicited beta power did not predict either positive (−0.05 < median*_diff_*s < 0.12, all HDIs overlapped with 0) or aversive (−0.06 < median*_diff_*s < 0.06, all HDIs overlapped with 0) memories. Moreover, there was no significant difference between the estimated trend of theta’s predictive capacity for positive and aversive memory retrieval (median*_diff_*s < 0.03, all HDIs overlapped with 0).

**Figure 6.**
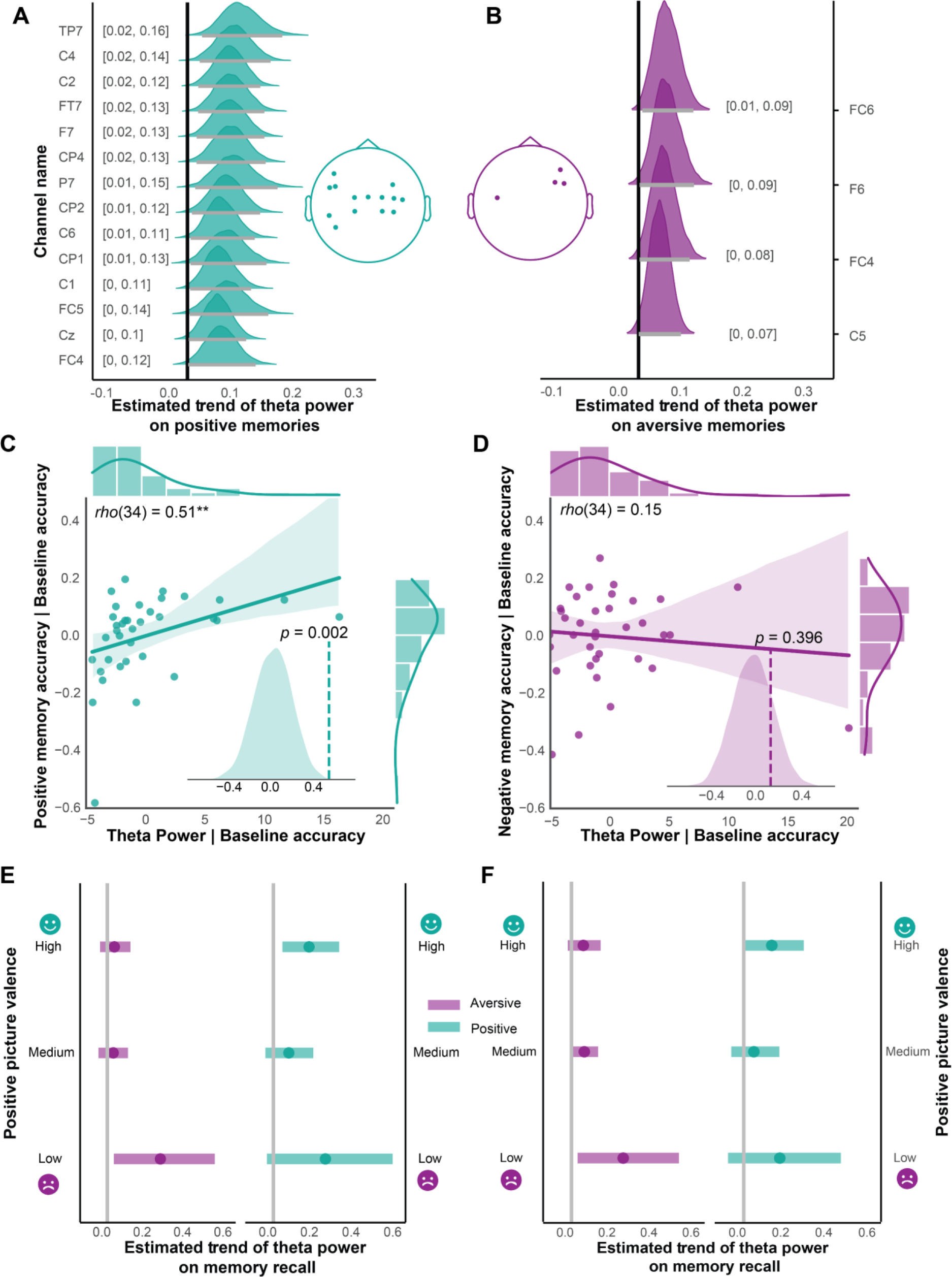
Prediction of TMR benefits through cue-elicited theta activity during NREM sleep at item- and participant-level. BLMM results of single-item theta power predicting positive (A) and aversive (B) memory recall accuracy across different channels. The x-axis represents the posterior distribution of the estimated trend, while the y-axis shows significant channels. The numbers in square brackets indicate the 95% HDI. Horizontal black lines represent the 95% HDI, and vertical black lines correspond to 0. If 0 does not fall within the 95% HDI, the result is considered significant. Topography illustrates significant channel locations. (C, D) Partial Spearman correlation of theta power (averaged across significant channels) with positive and aversive memory accuracy, while controlling for baseline memory accuracy. The inner density plot represents the distribution of permuted partial correlations, obtained by shuffling the relationship between power and memory accuracy 5000 times. The x-axis denotes the partial correlation values, and the dashed line indicates the actual partial correlation. (E, F) BLMM results of single-item theta power predicting positive memories, modulated by the affective rating of positive image valence. Theta power was averaged from the significant channels shown in Figures A and B. The Y-axis depicts the different levels of positive image ratings: low, medium, and high. Vertical gray lines indicate the value of 0, and horizontal lines represent the 95% HDI. Dots signify the median values.

To confirm the robustness of the results, we further examined this EEG-behaivoral relationship at the participant-level. We similarly found that cue-elicited theta power significantly predicted positive memory accuracy across most scalp electrodes (ps < 0.05, FDR corrected, Figure 6 C). However, no such relationship was observed for aversive memory (ps > 0.05, FDR corrected, Figure 6 D). Moreover, the correlation of theta power on positive memory accuracy was significantly stronger than that on aversive memory accuracy (zs > 2.13, p_corrected_s < 0.033). This finding suggested that at a participant-level, the cue-elicited theta power preferentially predicted positive over aversive memories.

We further investigated the impact of the positivity strength of the positive interfering memories (i.e., valence ratings of positive images) on the prediction of theta power for positive and aversive memories. Based on valence ratings of positive images (1 represents most aversive and 9 represents most positive), we categorized the positivity strength into low (1-3); medium (4-6) and high (7-9). We then employed a BLMM to explore how the positivity strength would modulate the prediction of theta on positive and aversive memories at an item level.

In this analysis, we included theta power (averaging across channels that showed significant theta prediction on positive memories), valence (aversive vs. positive memories), and positivity strength (low, medium, high), while accounting for time as covariates. Results revealed that when positivity strength was low, averaged theta power predicted aversive memories (median*_diff_* = 0.26, 95%HDI[0.03, 0.53]) but not positive memories (median*_diff_* = 0.25, 95%HDI[−0.03, 0.57]). Conversely, when positivity strength was high, averaged theta power predicted positive memories (median*_diff_* = 0.17, 95%HDI [0.04, 0.32]) but not aversive memories (median*_diff_* = 0.03, 95%HDI[−0.04, 0.11]). For medium positivity strength, average theta power did not predict neither aversive nor positive memories (median*_diff_*s < 0.07, all HDIs overlapped with 0, Figure 6E).

We next repeated the same analysis, but using theta power in predicting aversive memories (averaging across channels that showed significant theta prediction on aversive memories). The results indicated that when positivity strength was low, averaged theta power predicted aversive memories (median*_diff_* = 0.25, 95%HDI[0.03, 0.51])) but not positive memories (median*_diff_* = 0.17, 95%HDI[−0.07, 0.45]). For medium and high positivity strength, averaged theta power did not predict neither aversive nor positive memories (median*_diff_*s < 0.13, all HDIs overlapped with 0, Figure 6F).

Together, these findings suggested that cueing interfering memories preferentially reactivated the newly acquired positive memories rather than the older aversive memories. Importantly, when positivity strength was high, the positive interference effect was strong, with cue-elicited theta power preferentially predicting positive memories. When positivity strength was low, the positive interference effect was weak, with cue-elicited theta power preferentially predicting aversive memories. However, it shall be noted that the proportion of low positivity strength trials was small (low: 5.8%, medium: 57.6%, high: 36.6%). This helps explain that cue-elicited theta power predicted aversive memories only at the item level, but not at the participant level. Results with the delta bands showed a similar pattern to theta bands, while we did not observe any effect at sigma or beta bands (Figure S2).

In the noninterference condition wherein cues were only associated with aversive memories, we found that cue-elicited theta, sigma and beta power positively predicted aversive memories across multiple channels at the item-level (theta: median*_diff_*s > 0.07; sigma: median*_diff_*s > 0.08; beta: median*_diff_*s > 0.14), but not at the participant level (theta: rs < 0.33, ps > 0.64; sigma: rs < 0.24, ps > 0.97; beta: rs < 0.42, ps > 0.14, corrected). No effects were found for cue-elicited delta power at either item (median*_diff_*s < 0.037, all HDIs overlapped with 0) nor participant levels (rs < 0.34, p_corrected_s > 0.63, Figure S3).

### REM sleep modulated the TMR benefits on positive intrusions

Considering the influence of REM sleep on TMR and emotional memory (Cabrera et al., 2024; Batterink et al., 2017; Goldstein & Walker, 2014; Groch et al., 2013; Tamminen et al., 2017), we explored the relationship between REM parameters (percentage and REM-theta power) and behavioral measures (recall, intrusion, and positivity change scores). Consistent with previous analyses, we first combined the behavioral outcomes from the immediate and delayed tests. Spearman correlations in cued and uncued conditions did not find significant correlations after correction (p_corrected_s > 0.12; see Tables S5 and 6 for details).

Next, we examined the immediate and delayed tests separately. In the cued condition, we found that the REM percentage of Day 2 TMR night positively predicted positive intrusions during the delayed test (rho (33) = 0.45, p_corrected_ = 0.04) but not during the immediate test (rho (33) = 0.17, p_corrected_ = 0.33). Additionally, REM percentage did not predict aversive intrusions during either the immediate or delayed tests (p_corrected_s > 0.33, see Tables S7 and 8). In the uncued condition, no significant correlations were found (p_corrected_s > 0.50, see Tables S5-9). Furthermore, the correlation between the REM percentage from Day 2 sleep and delayed positive intrusion in the cued condition was significantly larger than in the uncued condition (z = 2.25, p = 0.01). These results suggest that the TMR effect on positive intrusions during the delayed test is contingent upon REM sleep following the NREM TMR.

## Discussion

This experiment provided new evidence that older aversive memories can be weakened via sleep reactivation of their corresponding positive memories that produced interference. Cue presentations during sleep also increased intrusions of positive memories during aversive memory recall. Moreover, cueing improved positive memories when earlier remembered aversive memories interfered with recall of positive memories during wakefulness. Regarding affect changes, TMR increased positive judgments toward memory cues, facilitating evidence accumulation toward these positive judgments. Sleep EEG results indicated that cue-elicited theta power predicted positive memories both at the item- and participant-levels, suggesting that cueing preferentially reactivated the recently acquired positive memories. Overall, our findings may offer new insights relevant for the treatment of pathological or trauma-related remembering.

Over the past decade, using TMR to modify memories during sleep has garnered much attention, particularly with fear conditioning and emotional episodes (Ai et al., 2015; Cairney et al., 2014; Hauner et al., 2013; He et al., 2015; Hutchison et al., 2021; Xia et al., 2023b). However, most of these studies used sleep cues associated with negative memories, yielding mixed results. We recently demonstrated that pairing a positive word with an aversive memory cue during NREM sleep, affect judgments became less negative (Xia et al., 2023b). Relatedly, research reported that recalling positive memories during wakefulness could reduce negative affect and ameliorate depressive symptoms (Speer et al., 2021; Speer & Delgado, 2017). We thus employed a novel approach to introduce positive interference to earlier acquired aversive memories during wakefulness, followed by reactivating these memories during NREM sleep. In line with our hypotheses, TMR not only weakened subsequent recall of these older aversive memories but also increased intrusions of positive memories. Intriguingly, we found that when remembered aversive memories interfered with the recall of positive interfering memories (i.e., those aversive remembered and positive forgotten items), TMR improved recall of these positive memories while weakening corresponding aversive memories. Our results were aligned with recent TMR research showing that episodic forgetting could be induced via reactivating interfering memories during sleep (Antony et al., 2018; Joensen et al., 2022; Oyarzún et al., 2017). Going beyond prior research on neutral memories, our results suggest that TMR preferentially reactivated recently acquired positive memories and weakened older aversive memories, thus altering the fate of emotional experiences.

One important goal of emotional memory editing is to alleviate the affective responses elicited by these memories. While TMR can be effective in modifying memories, subjective valence or arousal ratings might not be sensitive to TMR (Hu et al., 2020, but see Hutchison et al., 2021). Previous research suggested that a speeded affect judgment task might be better for capturing affective changes due to manipulation during sleep (Xia, et al., 2023b, see also Züst et al., 2019). Here, using a similar speeded binary affect judgment task, we found that TMR enhanced positive response changes in comparison to the uncued condition. Notably, our HDDM analyses showed that TMR selectively facilitated evidence accumulation toward the positive response, as evidenced by a higher drift rate. Existing studies have shown that the drift rate represents the quality of information obtained from stimuli during the evidence accumulation process, with a higher drift rate indicating lower random noise (Ratcliff et al., 2016; Ratcliff & McKoon, 2008), and memory can guide this process in preferential choice tasks (Gluth et al., 2015). Our results are also consistent with a previous TMR study in which TMR accelerated the evidence accumulation in a value-based binary choice task (Ai et al., 2018). Together, these results suggested that in addition to weakening aversive memories, TMR can also increase positive affective responses.

Complementing TMR-induced memory and affect benefits, our sleep EEG analyses provided further insights into the memory reactivation processes during sleep. First, compared to control sounds that were not paired with an image, we found that replaying memory cues elicited theta and beta EEG responses that may indicate memory reactivation (Creery et al., 2015; Schreiner et al., 2018; Schreiner & Rasch, 2015), with theta activity specifically linked to emotional memory reactivation (Canales-Johnson et al., 2019; Lehmann et al., 2016; Nishida et al., 2009; Xia, et al., 2023b). Second, examining EEG-behavioral relationships at the item level revealed that cued-elicited theta activity predicted both positive and aversive recall, while at the participant level, they predicted positive but not aversive recall. Moreover, when positivity strength was low (i.e., valence ratings were negative), cue-elicited theta power preferentially predicted aversive memories but not positive memories. However, when positivity strength was high (i.e., valence ratings were highly positive), cue-elicited theta power preferentially predicted positive memories. An important caveat is that only around 6% of positive images were rated as relatively aversive. These findings suggest that while TMR preferentially reactivated positive memories, aversive memories may be reactivated when their positive interfering memories were less positive. This observation can explain why we detected theta power’s prediction of aversive memories at the item level but not at the participant level.

Active system consolidation posits that newly acquired memories are temporarily stored in the hippocampus, and gradually become more cortical-dependent through the triple coupling of hippocampal ripples, thalamocortical spindles, and neocortical slow oscillations during NREM sleep (Diekelmann & Born, 2010; Klinzing et al., 2019; Paller et al., 2021). While most TMR studies have investigated memories acquired within hours before sleep (Hu et al., 2020), some results suggest that TMR may be less effective when reactivating 24-hour older memories (Seibold et al., 2018). Relatedly, using TMR to selectively reactivate overlapping memories can weaken older memories (acquired 3 hours ago) while strengthening recent contiguous learning (acquired 5 minutes ago), again highlighting that TMR may prioritize recent over earlier memories (Oyarzún et al., 2017). While these findings help explain why our TMR procedure preferentially reactivated recently acquired positive memories, an intriguing question remains — would TMR reactivate positive memories if they were introduced prior to the aversive memories. Future research is warranted to test whether TMR prioritizes recency over valence or vice versa. Furthermore, despite the lack of TMR effects from reactivating older aversive memories, we observed intriguing sleep EEG results: cues for older aversive memories also elicited greater delta-theta and sigma-beta power increases compared to control sounds. Importantly, cue-elicited theta power increases positively predicted post-TMR aversive memory accuracy at the item level. These findings suggested that even if older memories do not benefit from TMR, memory cues continue to be processed during sleep, as evidenced by changes in cue-elicited EEG activity (Cairney et al., 2018; Schreiner et al., 2018; Schreiner & Rasch, 2015; Wang et al., 2019).

Limitations and future directions should be noted. First, although our experiment aims to weaken aversive memories, the lab-induced emotional experiences of viewing aversive/positive images may not mimic typical traumatic experiences. Moreover, finding positive components within some highly traumatic experiences can be challenging. The generalizability of the current findings to clinical settings thus remains a critical goal for future research. Future research should explore ways to introduce positive interfering memories (Bjork et al., 1998), such as positive autobiographical memories or therapy-related memories (Speer et al., 2021; Schwartz et al., 2022), to effectively weaken real-life trauma memories. Second, the role of REM sleep in modulating emotional memories shall be further investigated (Cabrera et al., 2024; Goldstein & Walker, 2014). Our TMR was administered during SWS, a sleep stage that is critical for memory reactivation and systems consolidation (Klinzing et al., 2019; Hu et al., 2020; Wilson et al., 1994). Intriguingly, we also found that higher REM percentages during the TMR night predicted more TMR-induced positive memory intrusions, which is consistent with recent findings suggesting that the REM sleep could modulate the NREM TMR effects (Batterink et al., 2017; Pereira et al., 2022; Tamminen et al., 2017). Moreover, administering TMR during REM sleep can reduce subjective arousal of negative memories and the frequency of nightmares (Hutchison et al., 2021; Schwartz et al., 2022). Based on these promising results, future studies should further examine whether REM TMR could similarly weaken aversive memories.

Our results can also be related to retrieval-induced forgetting (RIF), in which repeated retrieval of selected memories would inhibit their related yet not retrieved memories, leading to forgetting of these non-retrieved memories(Anderson et al., 1994). Crucially, evidence suggests that selective retrieval would recruit the top-down inhibitory control to actively inhibit the interfering non-retrieved memory traces rather than disrupting specific reminder-memory associations (Wimber et al., 2015; Wessel & Anderson, 2024). While our findings and others (Antony et al., 2018a; Oyarzún et al., 2017; Joensen et al., 2022; Simon et al., 2018) suggest that TMR could induce forgetting during sleep, it remains unclear whether weakened memories would recover or generalize across reminders or contexts. Future studies should directly elucidate the underlying mechanisms and determine the longevity of the observed benefits.

In sum, our study presents a new approach for weakening older aversive memories during sleep. Notably, this benefit was achieved by introducing positive interference, followed by memory cueing during sleep. Cue-elicited theta power predicted the TMR benefit in recalling positive memories. More importantly, benefits were multifaceted, evidenced on voluntary recall, involuntary intrusions and speeded affect judgments. By demonstrating the memory and affect benefits of reactivating positive interfering memories, our study invites future research to harness the potential of sleep-based memory editing techniques in managing aversive memories, and promoting psychological well-being.

## Data availability

Pre-processed behavioral and EEG data are available on the Open Science Framework (OSF) upon publication: https://osf.io/servz/

## Code availability

Behavioral and EEG analysis codes are available on the Open Science Framework (OSF) upon publication: https://osf.io/servz/

## Materials and Methods

### Participants

A total of 37 participants (25 females and 12 males; Age: Mean ± S.D., 20 ± 2 years) were included in the behavioral analyses. For EEG analyses, we included 36 participants (one participant’s sleep EEG data was not saved properly due to an experimenter error). An additional 17 participants were recruited, but their data were excluded from subsequent analyses for the following reasons. Five participants’ data were dropped due to poor recognition accuracy (<50%) on Day 1 pre-sleep tests. Four participants were unable to fall asleep during the first night and one on the second night. Data from three participants were removed as they reported hearing the words while asleep. Data from four participants were excluded because they had fewer than 36 trials presented during sleep. Our sample size was determined based on recent within-subject TMR studies, with sample sizes ranging between 20 and 31 participants (Oyarzún et al., 2017; Schechtman et al., 2020; Schechtman et al., 2021; Hutchison et al., 2021; Antony et al., 2018b; Wang et al., 2019; Joensen et al., 2022; Ngo & Staresina, 2022).

All participants were native Chinese speakers, with self-reported regular sleep-wake cycles. They did not take any medications that influence sleep or mood. Furthermore, they had no history or current diagnosis of neurological or psychiatric illnesses. Participants were compensated with monetary incentives for their participation. The study received ethical approval from the Human Research Ethics Committee of the University of Hong Kong, HREC reference number: EA1904004. All participants provided written informed consent before participating in the experiment.

### Stimuli

We generated a set of 48 two-character pseudowords by randomly combining two neutral characters from the Chinese Affective Words System (Wang et al., 2008). The spoken words, which were used as auditory memory cues in later TMR, were produced using the Text-To-Speech function of iFLYTEK (duration: Mean ± S.D., 779 ± 56 ms). Visual stimuli consisted of images that were selected from four categories: animals, babies, people, and scenes. Each category included 12 positive and 12 aversive images, resulting in a total of 48 positive and 48 aversive images. These images were chosen from the International Affective Image System (IAPS; Bradley & Lang, 2017), the Nencki Affective Image System (NAPS; Marchewka et al., 2014), and various internet resources.

### Procedure

The experimental procedures were introduced to the participants prior to their first laboratory visit. Participants were told that the study included two consecutive nights of sleep in the laboratory, along with a series of computer tasks. As decribed below, these tasks were conducted during the daytime and nighttime with EEG recordings.

On Day 1 evening, participants arrived at the laboratory around 20:00. After EEG setup, they completed the following tasks in order: 1) aversive image rating, 2) cue (pseudowords) - target (aversive image) encoding (48 trials in each of 2 blocks), 3) cued recall of aversive images (2 blocks), 4) cue (pseudowords) - target (aversive image) encoding (2 blocks), 5) cued recall of aversive images (2 blocks). The first night serves as an adaptation night, facilitating the consolidation of the newly learned associations. Participants went to bed around 23:00, with EEG recorded throughout the night.

On Day 2 morning, participants were awakened at around 7:00. After breakfast and freshening up, they began the post-sleep tasks at approximately 7:30, which included: 1) a speeded affect judgment task; 2) affect ratings; 3) cued recall of aversive images. Participants left the lab upon completing these tasks.

On Day 2 evening, participants returned to the lab at around 20:30. Following EEG setup, they completed the following tasks in order: 1) positive image rating; 2) cue (pseudoword) - target (positive image) encoding (2 blocks); 3) cued recall (2 blocks); 4) cue (pseudoword) - target (positive image) encoding (2 blocks); 5) cued recall (2 blocks). Participants went to bed around 23:00, and half of the spoken words were played as memory cues during NREM sleep until approximately 02:00.

On Day 3 morning, participants were provided breakfast upon waking, and then completed the following tasks in order: 1) a speeded affect judgment task; 2) affect ratings; 3) cued recall of positive memories with self-report of involuntary intrusions of aversive memories; and cued recall of aversive memories with self-report of involuntary intrusions of positive memories. The order of positive and aversive cued recall tasks was counterbalanced across participants.

On Day 5, participants returned to the lab around 9:00 and completed the same tasks as on Day 3 morning: 1) a speeded affect judgment task; 2) affect ratings; 3) cued recall of positive or aversive memories with self-report of involuntary intrusions (the order of positive and aversive recall tasks was counterbalanced across participants). Lastly, participants gave semantic similarity ratings of the aversive and positive images.

### Tasks

#### Aversive (Day 1 evening) and positive (Day 2 evening) encoding

During the Day 1 evening aversive memory encoding task, participants learned 48 unique cue-target pairs involving pseudowords (cues) and aversive images (targets). These 48 images consisted of 12 images from each of four content categories (animals, babies, people, and scenes). During the Day 2 evening session, we randomly selected cues from half of the pairs (i.e., 24 word-image pairs, with images from two out of the four categories) and paired these cues with positive images selected from the same two categories, thus introducing positive interfering memories. This design created two conditions: the interference condition (pseudowords associated with both an aversive image on Day 1 and a positive image on Day 2) and the non-interference condition (pseudowords associated with only an aversive image on Day 1).

The encoding session included four blocks, each containing a passive viewing task and a recognition task with feedback. During the passive viewing task, each trial began with fixation (800-1200 ms), followed by simultaneous auditory and visual presentation of cues (approximately 1000 ms). Subsequently, auditory and visual cues were displayed again along with the image (2000 ms). Visual cues were shown in the top half of the screen, while the image was shown in the center. After all pairs (48 word-image pairs for Day 1 aversive encoding and 24 word-image pairs for Day 2 positive encoding), participants took a 1-minute break before starting the recognition task with feedback.

During the recognition task, each trial began with fixation (800-1200 ms), followed by simultaneous auditory and visual presentation of cues (1500ms), along with three images displayed in the center of the screen. The correctly paired image was randomly presented in one of three locations. The two filler images were selected from other learned word-image pairs. Participants used keys “1”, “2”, or “3” to indicate the image that was paired with the cue. Regardless of the participant’s response, the correct cue-target pair was presented aurally and visually again for 1000 ms as feedback. After this recognition with feedback task, the mean recognition accuracy was displayed in the center of the screen for 3000 ms. To increase encoding accuracy, participants completed a cued verbal recall task after two blocks of passive viewing and recognition with feedback tasks, followed by another two blocks of passive viewing and recognition with feedback tasks. Subsequently, participants completed the cued verbal recall task again, and the memory performance from this task was used in subsequent analysis.

#### Image/cue/word rating task

In the image-rating task, participants rated 48 aversive and positive images using a 9-point Likert scale along valence (extremely aversive to extremely positive) and arousal (extremely calm to extremely excited). Each trial began with an 800-ms fixation, followed by the presentation of images at the center of the screen. After participants submitted their ratings using a computer mouse, a 500-ms blank screen was displayed. Images were presented in a random order. Results indicated that individuals consistently perceived positive images as positive and negative images as negative (aversive: 2.99 ± 0.10; positive: 7.03 ± 0.12, t(36) = 22.0, p < 0.001, **d** = 3.61).

In the cue-affect-rating task, participants assessed the valence and arousal of pseudowords while hearing them spoken. The procedure was otherwise identical to that of the image-rating task.

#### Speeded Affect-judgement task

In the speeded affect-judgment task (Day 2, Day 3, and Day 5 mornings), each trial began with an 800 −1200ms fixation, followed by simultaneous auditory and visual presentation of cues at the center of the screen for approximately 1000 ms. Participants were instructed to make a rapid binary judgment, endorsing the cue as either aversive or positive, by pressing the left or right keys within 1.5s. The inter-trial interval was 1s. Participants completed four blocks of the task (192 trials). The results from the four blocks were subsequently averaged per cue to obtain participants’ affect responses towards the memory cues.

#### Cued recall and involuntary intrusion task

In the cued recall task, each trial began with a fixation period lasting between 800 and 1200 ms across each trial, followed by the aural presentation of the cue for approximately 1000 ms. Participants were instructed to provide a detailed verbal description of the images paired with the cue within a 15-s interval when the word “speaking” appeared on the screen. A microphone was used to capture responses. The inter-trial interval of 3 seconds.

Two independent raters listened to memory responses to determine if the image was uniquely and correctly described (see also Catarino et al., 2015). In any case of inconsistent ratings, a third rater reconciled the discrepancy. All raters were blind to the experimental conditions.

In the aversive cued recall task (Day 3 and Day 5 morning), participants were also instructed to report any involuntary intrusion of positive memories on each trial. They were specifically asked to indicate whether a positive memory intrusion occurred while recalling the aversive memory. Similarly, in the positive cued recall task (Day 2 evening and Day 3 and Day5 mornings), participants were instructed to report any involuntary intrusion of aversive memories on each trial. Participants rated the frequency of involuntary intrusions on a scale ranging from 0 to 2, where 0 represented “never,” 1 represented “rarely,” and 2 represented “often”. Ratings of 1 and 2 were combined to indicate the presence of intrusions, while a rating of 0 indicated no intrusion (Hellerstedt et al., 2016; Levy & Anderson, 2012).

#### Semantic similarity rating

In the final session on Day 5, participants assessed semantic similarities between aversive and positive images that were paired with the same cues. This evaluation aimed to determine the extent of interference, as images with higher conceptual similarity but opposite valence may engender stronger interference (Antony & Bennion, 2023; Goh & Goh, 2006; Rose & Abdel Rahman, 2017). Each trial commenced with a 500 ms fixation, followed by simultaneous side-by-side presentation of aversive and positive images that were paired with the same pseudoword (interference trials). Participants used the mouse to rate the semantic similarity between the two images using a 5-point Likert scale, ranging from 1 (not similar at all) to 5 (extremely similar). We examined whether semantic similarity rating between two images would influence TMR effect or EEG-behavioral relationship. Nevertheless, our study did not reveal any significant findings.

#### TMR setup

We randomly selected one category from the interference condition (12 pseudowords) and one category from the noninterference condition (12 pseudowords) to serve as cued items in the TMR procedure. The remaining two categories served as uncued items from interference and noninterference conditions. Additionally, 12 novel pseudowords that were never paired with any images were introduced as control sounds, as they would be unlikely to trigger any retrieval of the experimental stimuli. Consequently, a total of 36 pseudowords (interference, noninterference, control) were played during sleep. The 36 cues were presented randomly in blocks during NREM sleep. Trained experimenters initiated cueing when participants had been in SWS for a minimum of 5 minutes. Cueing was discontinued if participants entered REM or N1 sleep or exhibited arousal or wakefulness (i.e., bursts of muscle activity or alpha activity). Within each block, cues were presented with an inter-stimulus interval (ISI) of 4 seconds, with each cue lasting for 1 second, resulting in a 3-minute block duration. Each block was separated by a 60-second interval. The cueing process was terminated upon reaching either (1) 20 blocks of cues, amounting to a total of 720 trials, or (2) 02:00, whichever occurred first.

All experimental tasks were carried out using Psychopy 3.0 software (Peirce, 2007). During TMR, auditory cues (i.e., spoken words) were delivered at an approximate sound pressure level of 38 dB through a loudspeaker situated approximately 50 centimeters above the bed. White noise was played as background throughout the entire night.

#### EEG recording and preprocessing

Sleep was monitored and recorded using high-density electroencephalography (EEG), electrooculography (EOG) for eye movement recording, and mentalis electromyography (EMG) for muscle activity recording. EEG data were collected using a 64-channel system (eego, eemagine, ANT, The Netherlands), and impedances were maintained below 20 kΩ. Signals were sampled at 500 Hz online, using CPz as the reference. Prior to sleep, two EMG electrodes were attached bilaterally to the mentalis regions, with a ground electrode placed on the forehead. For monitoring only, EEG data were bandpass filtered at 0.5 Hz to 40 Hz, and EOG and EMG were not filtered.

All EEG processing steps were carried out using MNE-Python (Gramfort, 2013) and Python 3.6. The processing steps included the following: (1) down-sampling the EEG data to 200 Hz, (2) applying a 50 Hz notch filter along with 0.1 - 40 Hz bandpass finite impulse response (FIR) filters, (3) visually identifying and marking bad channels, (4) re-referencing the data to the average of all non-marked electrodes after excluding M1 and M2, and (5) segmenting continuous EEG data into long epochs (20s epochs ranging from −10 to 10 seconds relative to cue onset) for subsequent time-frequency analysis and sleep event detection. Finally, artifact epochs were removed through visual inspection, followed by interpolation of bad channels.

#### Behavioral analysis

For the speeded affect judgment task, we computed the positive response ratio, which ranged from 0 (all responses were aversive) to 1 (all responses were positive). We then calculated the affect judgment change score by subtracting the pre-sleep positive response ratio from the post-sleep immediate and delay ratios, resulting in a range from −1 to 1, with a larger score indicating increased positive responses from pre-sleep to post-immediate or post-delay, and a zero score indicating no changes in affect judgments.

For the affect rating task, we calculated affect rating changes by subtracting pre-sleep baseline valence/arousal ratings from post-sleep immediate and delay ratings. A higher valence/arousal change score indicated a more positive or arousing change from pre-sleep to post-sleep immediate or delay. We then submitted the affect judgment change, valence change, and arousal changes to a Bayesian model to quantify the TMR effect at an item level. Regarding memory changes, we employed Bayesian logistic regression to quantify the TMR effect on memory accuracy and memory intrusion at the item level, given the binary nature of the data (i.e., recalled or not, experienced intrusion or not).

#### EEG Time-Frequency Analyses

In the time-frequency analysis, the initial step was cropping epochs to the interval −2 to 6 s relative to TMR onset. Next, a continuous wavelet transformation featuring variable cycles (3 cycles in length at 1 Hz, increasing linearly with frequency up to 15 cycles at 40 Hz) was applied to the cropped epochs. We thus extracted power values from 1 to 40 Hz in increments of 0.5 Hz and 5 ms intervals. In order to mitigate edge artifacts, epochs were further cropped to −1 to 4 s relative to TMR onset. Subsequently, spectral power normalization was applied using Z-score transformation of all trials, with a baseline interval from −1 to −0.2 s. Trial-level time-frequency data were retained for further analysis. For the contrast between memory cues and control sounds, trials were averaged for each specific experimental condition within individual participants. A three-dimensional spatial-temporal-frequency permutation test (two-tailed, one-sample with randomization of 1000 and a statistical threshold of 0.05) was utilized to evaluate significant clusters. This method allowed for three comparisons: interference and control sound conditions, noninterference and control sound conditions, and interference and noninterference conditions.

#### Sleep staging

Sleep stage analyses were conducted using a machine learning algorithm, the Yet Another Spindle Algorithm (YASA, Vallat & Walker, 2021). Staging results were next validated by an experienced sleep researcher. In adherence to YASA’s recommendations, EEG data were initially re-referenced to FPz (or Fp2 if Fpz was labeled as a bad channel). The C4 (or C3 in instances where C4 was labeled as a bad channel), and EOG served as input for the algorithm. Five participants on Day 1 and one participant on Day 2 had Ground or Reference channels disconnected during the second half of the night due to head and body movements, resulting in signal loss of all channels. As a result, we excluded these participants when reporting sleep staging information. Before calculating sleep staging statistics, artifacts were identified and removed. TMR cueing accuracy was determined offline based on the results of this automatic sleep staging. Further details on sleep stages and cueing accuracy can be found in Table S4.

#### Drift diffusion modeling

The Drift Diffusion Model (DDM) was used to decompose the decision-making processes, which assesses sensory inputs and internal processing related to binary choices by continuously combining sensory information related to two distinct choices (positive and aversive responses). This integration proceeds until a certain threshold is achieved, signifying that a sufficient amount of evidence has been collected to confidently make a decision (Myers et al., 2022). The DDM separates the decision process into four key parameters: the starting point (z) between the two choice boundaries, which reflects a pre-decision bias; the non-decision time (t), which accounts for decision-irrelevant factors; the noisy drift process (v), which represents the accumulation of information toward the preferred choice boundary; and the choice boundaries (a), which mark the completion of the decision-making process when the accumulated evidence reaches either one of them. We utilized a Bayesian hierarchical estimation of the Drift Diffusion Model (HDDM 0.8) implemented in the Docker HDDM framework (Pan et al., 2022). Our hierarchical design considered that individual participant parameters are not entirely independent but are drawn from a shared distribution. We hypothesized that both drift rate (v) and starting point (z) would be influenced by TMR manipulation and test time in the interference and non-interference conditions. Additionally, we allowed the choice boundary (a) and non-decision time (t) to vary based on TMR and test time. To approximate the posterior distribution of the model parameters, we employed Markov Chain Monte Carlo (MCMC) sampling, generating 25,000 samples and discarding 2,000 as burn-in. Model convergence was assessed through visual inspection of the traces and autocorrelation of the model parameters, as well as by computing the Gelman-Rubin R-hat statistic (R-hat<1.1, Gelman et al., 2014).

Due to our assumption that only drift rate and pre-decision bias would be affected by our experimental condition, we focused on these parameters. We applied HDDM regression analysis using the following model:

[1] v = 1 + TMR + Time +TMR*Time
[2] z = 1 + TMR + Time +TMR*Time
[3] v, z = 1 + TMR + Time +TMR*Time

In the above models, the decision boundary (a) and non-decision time (t) were estimated at the individual level. To identify the best-fitting model, we conducted model comparisons using leave-one-out cross-validation of the posterior log-likelihood (LOO-CV) as implemented in the ArviZ package for Python (Kumar et al., 2019). The top-ranked model, determined by “arviz.compare”, was v = 1 + TMR + Time + TMR*Time, and thus was reported.

#### Statistics

In the analysis of behavioral data, we employed a paired sample t-test on all 48 memory items to examine the change in aversive memory from pre-sleep to post-sleep on Day 1.

Additionally, a repeated measures ANOVA was conducted to assess the interference memory performance across aversive pre-sleep (Day 1 evening), aversive post-sleep (Day 2 morning), and positive pre-sleep conditions (Day 2 evening). These analyses were conducted to examine whether participants successfully encoded aversive memories, whether aversive memories were consolidated following sleep, and how prior aversive memories influenced later encoding of positive memories when they shared the same cues.

To measure TMR effects, we employed the BLMM to analyze the effect of TMR in the interference and non-interference conditions. To determine the best-fitted model, model comparisons were performed utilizing leave-one-out cross-validation of the posterior log-likelihood (LOO-CV) combined with Pareto-smoothed importance sampling, which was implemented in the loo package for R (Vehtari et al., 2018).

Non-informative priors were utilized in all models. For each model, 4 Markov Chain Monte Carlo (MCMC) chains were executed using 5,000 samples, with the initial 500 samples discarded as a warm-up phase. The Gelman-Rubin r-hat statistic was employed to assess model convergence (R-hat < 1.1; Gelman & Rubin, 1992). Statistical inferences were based on the 95% highest density interval (HDI) of the posterior distribution. Effects were considered significant if the 95% HDI did not encompass 0.

#### EEG-behavioral correlation

In the time-frequency analysis, we employed a cluster-based two-tailed, one-sample permutation test across channels, time points, and frequency bands with 1000 randomizations and a statistical threshold of 0.05 against zero. An alpha level of 0.05 was used for statistical significance testing throughout the manuscript. For effect sizes, we reported Cohen’s dz in our within-subject design, accompanying paired sample t-tests.

We first investigated whether cue-elicited EEG power was associated with subsequent memory accuracies (remember vs. forgotten) for positive and for aversive memories. To achieve this, we employed the following BLMM:

> EEG power = 1 + memory accuracy * valence + time + (1 + memory accuracy * valence | participant)

This model allowed us to examine the association between cue-elicited EEG power and subsequent memory accuracy of different emotional valences, while accounting for individual participant variability.

To quantify the relationship between cue-elicited EEG power and behavioral measures, we utilized BLMM. EEG Power was extracted from significant clusters at the item level, and then the cue-elicited power values were submitted to BLMM to predict memory accuracy, memory intrusion, and positive response changes. For interference condition, we were interested in the interaction between EEG power and valence (aversive vs. positive); thus, the following BLMM was employed:

> Memory accuracy = 1 + EEG power * valence + baseline accuracy + time + (1 + power*valence | participant)

By adding baseline accuracy to the model, we aimed to explore the predictive effect of cue-elicited power on TMR-induced memory changes. This analysis was repeated across all channels and for delta, theta, sigma, and beta bands.

For noninterference condition, the following BLMM was employed:

> Aversive memory = 1 + EEG power + baseline accuracy + time + (1 + power | participant)

To further investigate whether the affect rating of positive images modulated the power’s prediction on aversive and positive memories, we averaged the significant channels that predicted positive or aversive memories and submitted them to the following BLMM:

> Memory accuracy = 1 + EEG power * valence * categorical positive image valence + time + (1 + power * valence * categorical positive image valence | participant)

In this model, the categorical positive image valence represents the division of positive image valence ratings into three conditions (low, medium, high) using the ‘cut’ command in R. All other variables were retained from the previous model, while baseline accuracy was excluded to decrease model complexity.

For the analysis of EEG-behavioral relationships, we conducted each model across all channels. It is important to note that in Bayesian statistics, there is typically no requirement for multiple comparison corrections (Gelman et al., 2012; Kruschke, 2013). This characteristic of the Bayesian approach allows for a more straightforward interpretation of results across multiple channels in the context of our study.

## Acknowledgments

We would like to thank Dr. Menglu Chen for offering valuable feedback and suggestions on this work and the visualizations. We thank Dr. Chuanpeng Hu for his assistance with the HDDM Docker installation and insightful suggestions for the HDDM analysis. We thank Yuheng Shi, Lingqi Zhang, Ruoying Zheng, and Xibo Zuo for their help in data collection and memory coding. The research was supported by the Ministry of Science and Technology of China STI2030-Major Projects (No. 2022ZD0214100), National Natural Science Foundation of China (No. 32171056), the General Research Fund (No. 17614922) of Hong Kong Research Grants Council to X. H. The funders had no involvement in the study design, data collection and analysis, publication decisions, or manuscript preparation.

