## Supplementary material for "Aversive memories can be weakened during human sleep via the reactivation of positive interfering memories": supplmental Materials

**This PDF file includes:**

Figures S1 to S4
Tables S1 to S9
SI References

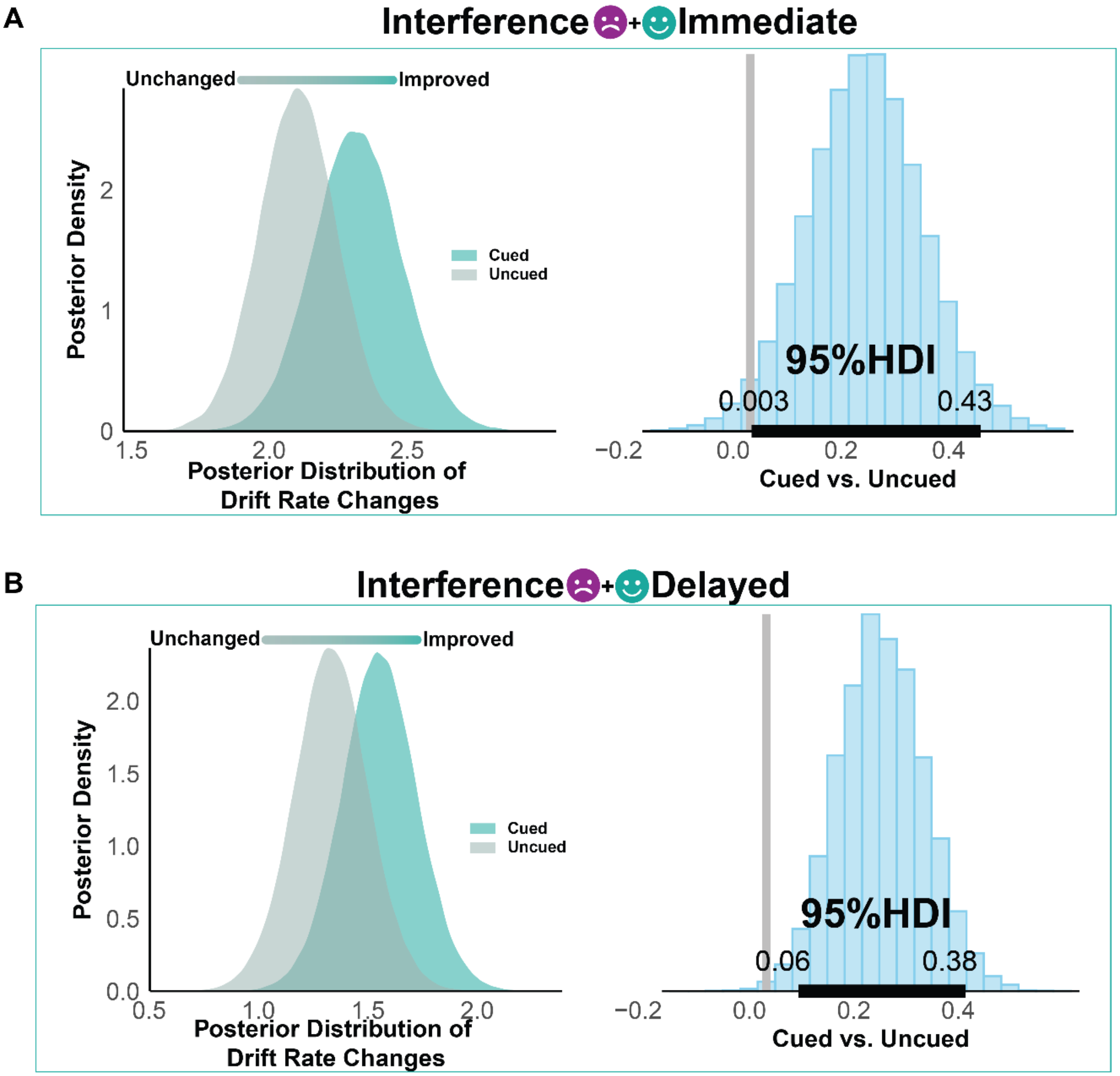


**Fig. S1. Immediate and delayed results of drift rate changes during speed choices task in the interference condifion** TMR facilitated evidence accumulation towards a positive response in the affect judgment task during immediate and delayed test. Drift rates were calculated using the HDDM regression module. Panels A and B present drift rate changes across cued and uncued conditions during immediate and delayed tests, respectively. In both panels, green squares signify a positive dependent measure, while the left panel density plots depict the posterior distribution of related dependent variables in cued and uncued conditions. The right panel histogram plots display the contrast between cued and uncued conditions. Horizontal black lines represent the 95% highest density interval (HDI), and vertical gray lines correspond to 0. Results are considered significant if 0 does not fall within the 95% HDI.

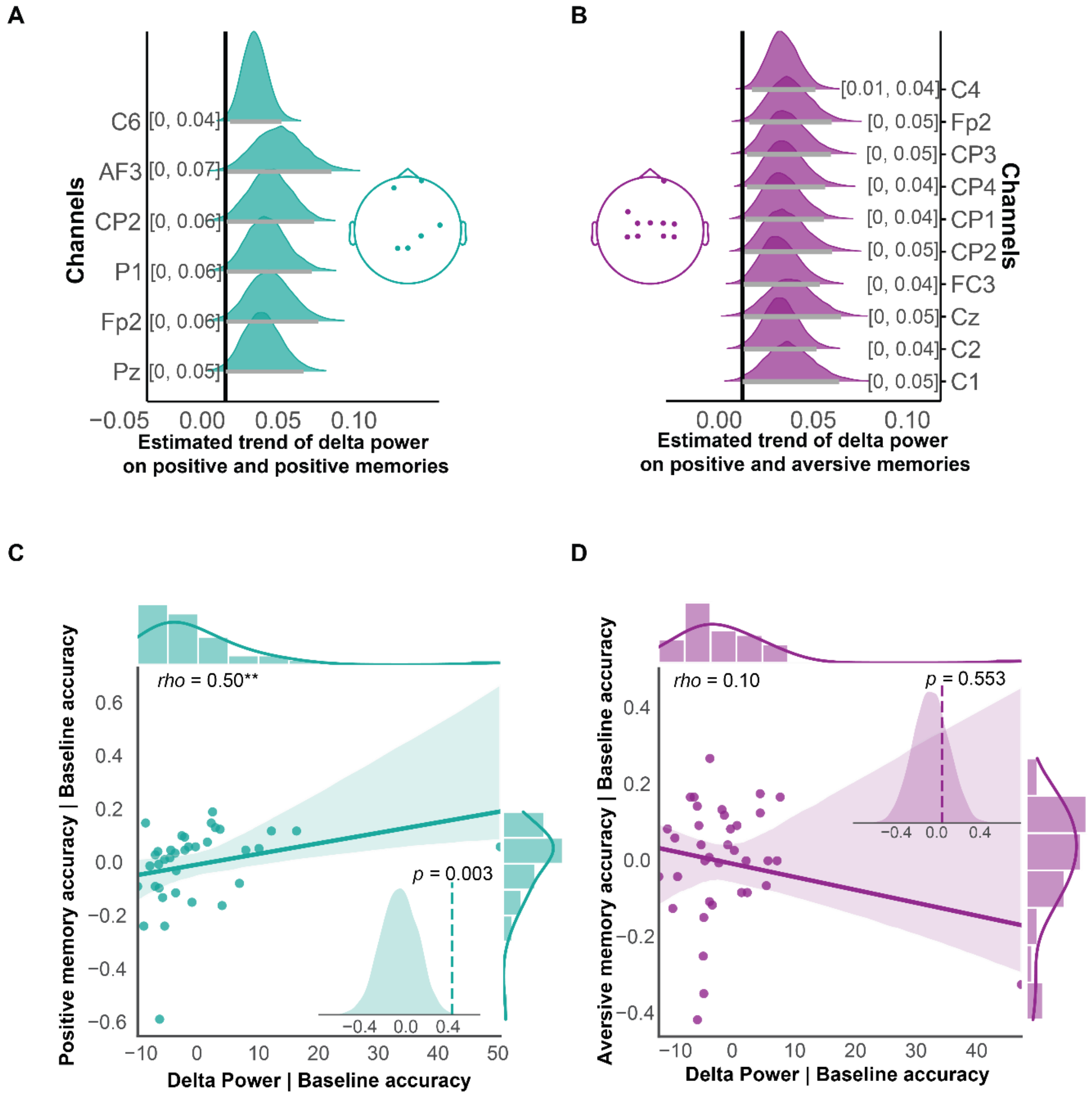


**Fig. S2. Prediction of TMR benefits through cue-elicited delta activity during NREM sleep at item- and subject-level in the interference condition**. BLMM results of single-item delta power predicting (A) positive and (B) aversive memory recall accuracy across different channels. The x-axis represents the posterior distribution of estimated trend, while the y-axis shows significant channels. The numbers in square brackets indicate the 95% HDI. Horizontal black lines represent the 95% HDI, and vertical gray lines correspond to 0. If 0 does not fall within the 95% HDI, the result is considered significant. (C, D) Partial Spearman correlation of delta power (averaged across significant channels) with positive and aversive memory accuracy, while controlling for baseline memory accuracy. The inner density plot represents the distribution of permuted partial correlations, obtained by shuffling the relationship between power and memory accuracy 5000 times. The x-axis denotes the partial correlation values, and the dashed line indicates the actual partial correlation.

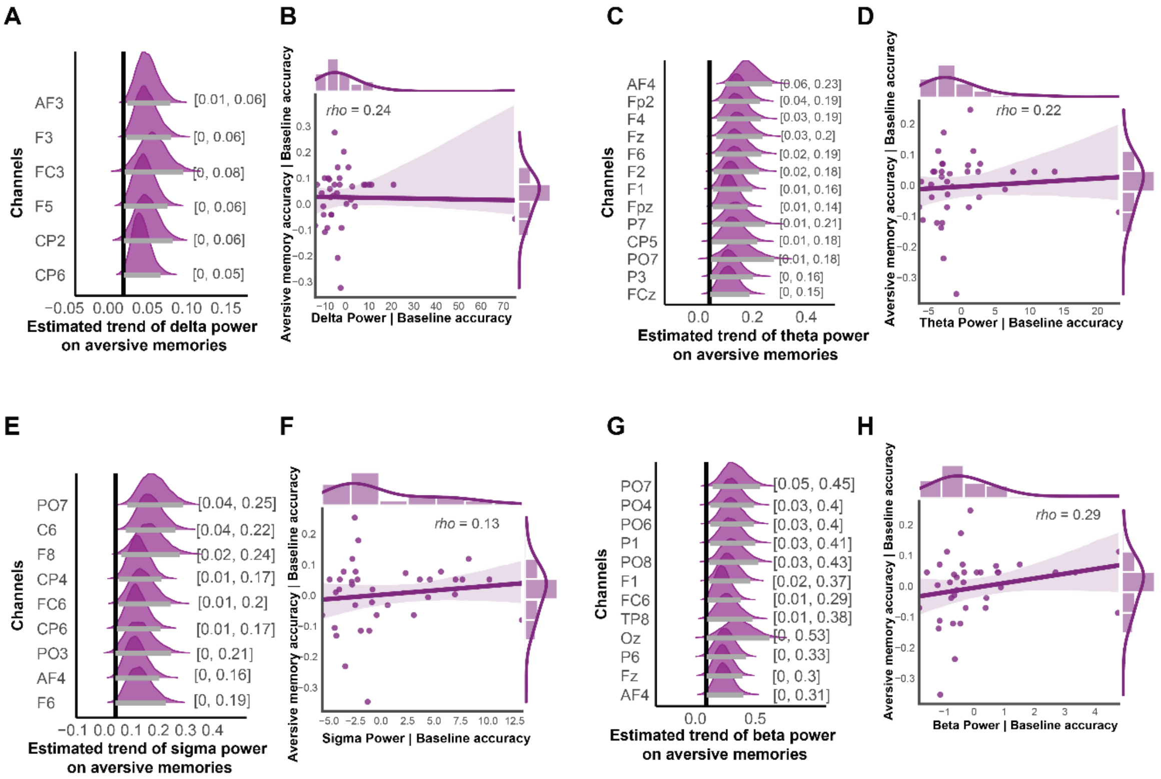


**Fig. S3. Prediction of TMR benefits through cue-elicited activity during NREM sleep at item- and subject-level in the noninterference condition.** (A, C, E, G) BLMM results of single-item (A) delta, (C) theta, (E) sigma, and (G) beta power predicting aversive memory recall across different channels. The x-axis represents the posterior distribution of estimated trend, while the y-axis shows significant channels. The numbers in square brackets indicate the 95% HDI. Horizontal gray lines represent the 95% HDI, and vertical black lines correspond to 0. If 0 does not fall within the 95% HDI, the result is considered significant. (B, D, F, H) Partial Spearman correlation of (B) delta, (D) theta, (F) sigma, and (H) beta power (averaged across significant channels) with aversive memory accuracy, while controlling for baseline memory accuracy.

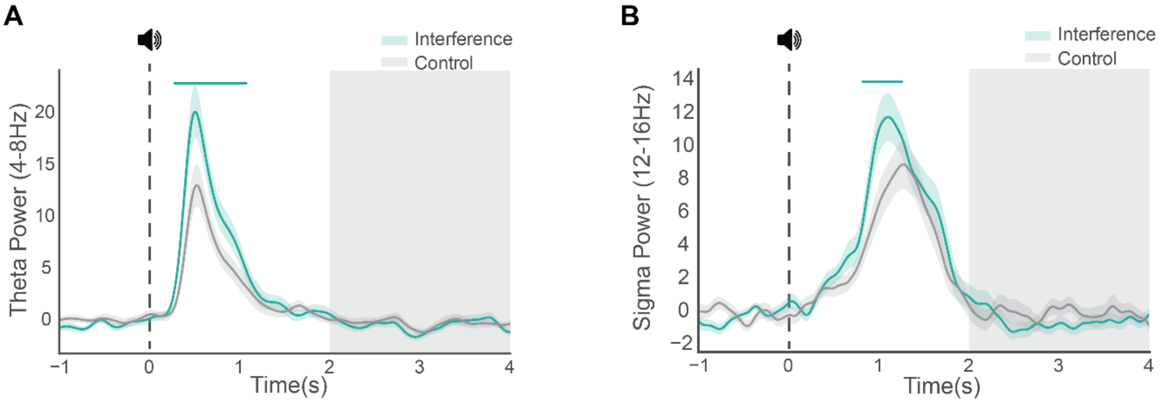


**Fig. S4.** This figure displays the differences in theta and sigma power between interference cues and control sounds at the Cz electrode, which was selected in accordance with previous research (Schechtman et al., 2021). Results remain consistent when using a cluster of fronto-central-parietal channels. Horizontal green lines indicate significant time windows. The vertical dashed line at 0 represents the onset of the auditory cue. Shaded areas denote the time windows used to examine EEG power reduction. Interference cues elicited significantly larger theta and sigma power than control sounds around 1 second, while no significant differences were observed between memory cues and control sounds within the 2-4 second range for both theta and sigma bands. To investigate the potential link between EEG power decreases during TMR and subsequent memory, we extracted cue-elicited time-frequency resolved theta and sigma power from the interference and control conditions. Consistent with our main findings (Figure 5AB), interference memory cues elicited a theta and sigma power increase around 1 second post-cue (pclusters = 0.001, indicated by the horizontal green line). Moreover, we observed that interference cues were associated with reduced sigma power relative to the control sounds during the 2-4 seconds post-cue (indicated by the gray shaded areas), though this reduction did not reach statistical significance ($t$ (35) = -1.59, $p$ = 0.122). Furthermore, theta power did not differ between the interference cues and the control sound during 2-4 seconds post-cue (see the figure below, $t$ (35) = -0.83, $p$ = 0.413). We proceeded to examine the relationship between theta and sigma power decreases and the TMR effect on memory accuracy. We averaged trial-level EEG power data from 2-4 s post-cue at Cz. We then employed a BLMM with EEG power (sigma or theta) and valence (positive vs. aversive memories) as fixed factors, incorporating baseline memory accuracy and time as covariates. The results did not reveal any significant predictions of sigma (-0.008 < mediandiffs < 0.015; all HDIs overlapped with 0) or theta (0.001 < mediandiffs < 0.099; all HDI overlapped with 0) power decrease on aversive or positive memory accuracy. In addition to using the signal from Cz, we also extracted trial-level power from the frontal-central-parietal area (Cz, C1, C2, CP1, CP2, F1, F2, Fz) according to previous studies**(1–3)**. The results remained consistent such that cue-elicited sigma or theta power decrease did not significantly predict negative or positive memories (all HDIs overlapped with 0).

**Table S1. Descriptive statistics of key dependent variables (Mean ± SE).** The 'Selected' represents the items remembered during the aversive memory baseline test but were forgotten during the positive memory baseline test.

| **Dependent variables** | **TMR** | **Interference** | **Immediate** | **Delayed** |
|---|---|---|---|---|
| Aversive memories | Cued | Interference | 0.768 ± 0.030 | 0.734 ± 0.036 |
| | | Noninterference | 0.860 ± 0.028 | 0.849 ± 0.028 |
| | Uncued | Interference | 0.860 ± 0.029 | 0.851 ± 0.031 |
| | | Noninterference | 0.813 ± 0.026 | 0.786 ± 0.028 |
| Positive intrusions | Cued | Interference | 0.822 ± 0.033 | 0.664 ± 0.044 |
| | | Noninterference | 0.070 ± 0.018 | 0.063 ± 0.017 |
| | Uncued | Interference | 0.788 ± 0.036 | 0.592 ± 0.044 |
| | | Noninterference | 0.065 ± 0.017 | 0.056 ± 0.014 |
| Positivity change socres | Cued | Interference | 0.562 ± 0.038 | 0.366 ± 0.040 |
| | | Noninterference | 0.014 ± 0.019 | -0.016 ± 0.020 |
| | Uncued | Interference | 0.508 ± 0.039 | 0.312 ± 0.042 |
| | | Noninterference | 0.003 ± 0.026 | -0.031 ± 0.021 |
| Positive memories | Cued | Interference | 0.899 ± 0.027 | 0.844 ± 0.028 |
| | Uncued | | 0.899 ± 0.021 | 0.829 ± 0.029 |
| Aversive memory accuracy (selected) | Cued | Interference | 0.656 ± 0.057 | 0.649 ± 0.062 |
| | Uncued | | 0.884 ± 0.034 | 0.788 ± 0.047 |
| Positive memory accuracy (selected) | Cued | Interference | 0.845 ± 0.047 | 0.817 ± 0.056 |
| | Uncued | | 0.761 ± 0.056 | 0.724 ± 0.058 |

**Table S2.** Summary of Bayesian Linear Mixed Models and Model Comparisons. This table summarizes all Bayesian linear mixed models referenced in the Results section, along with model comparisons for including time as a fixed factor or covariate, and corresponding references to figures and tables. To identify the best-fitting model, model comparisons were conducted using leave-one-out cross-validation of the posterior log-likelihood (LOO-CV), combined with Pareto-smoothed importance sampling, as implemented in the loo package for R (Vehtari et al., 2018). The optimal model was selected based on the highest expected log point-wise predictive density (ELPD). A value of 0 for ELPD and SE_DIFF in the table indicates superior model performance. However, a comparison is deemed significant only when the absolute mean difference in ELPD between any two models (elpd_diff in brms) surpasses twice the standard error of the differences (2*SE_diff). AM, aversive memories; BAM, baseline aversive memories; PM, positive memories; BPM, baseline positive memories; AI, aversive intrusions; BAI, baseline aversive intrusions; MR, memory recall; BM, baseline memory recall; PI, positive intrusions; PC, positivity change score; RT_change, reaction time change; VC, valence changes; AC, arousal changes.

| | Models | Model Comparasion | | Figure/Results |
|---|---|---|---|---|
| | | **Elpd_diff** | **SE_diff** | |
| #1 | AM = 1+TMR*Interference + time + BAM + (1 + TMR*Interference \| subject) | 0 | 0 | Figure 3 AB |
| #2 | AM = 1+TMR*Interference*time + BAM + (1 + TMR*Interference*time \| subject) | -5.1 | 1.8 | Table S3 |
| #3 | PM = 1+TMR+time +BPM + (1+TMR \| subject) | -0.1 | 2.1 | |
| #4 | PM = 1+TMR*time +BPM + (1+TMR*time \| subject) | 0 | 0 | Table S3 |
| #5 | AI = 1+TMR+time +BAI + (1+TMR \| subject) | -7.6 | 5.1 | |
| #6 | AI = 1+TMR*time +BAI + (1+TMR*time \| subject) | 0 | 0 | Table S3 |
| #7 | MR = 1 + TMR*valence + time + BM + (1 + TMR*valence \| subject) | -2.5 | 4.5 | |
| #8 | MR = 1 + TMR*valence*time +BM + (1+TMR*valence*time\| subject) | 0 | 0 | Table S3 |
| #9 | MR = 1 + TMR*valence + time + (1 + TMR*valence \| subject) | 0 | 0 | Figure 3 CD |
| #10 | MR = 1 + TMR*valence*time + (1+TMR*valence*time\| subject) | -5.6 | 2.3 | Table S3 |
| #11 | PI = 1 + TMR * Interference + time + (1+TMR*Interference \| subject) | -21.4 | 7.7 | Figure 3 EF |
| #12 | PI = 1 + TMR * Interference*time + (1+TMR*Interference*time \| subject) | 0 | 0 | Table S3 |
| #13 | PC = 1 + TMR * Interference + time + (1+TMR*Interference \| subject) | -33.6 | 10.1 | Figure 4 AB |
| #14 | PC = 1 + TMR * Interference*time + (1+TMR*Interference*time \| subject) | 0 | 0 | Table S3 |
| #15 | RT change = 1 + TMR * Interference + time + (1+TMR*Interference \| subject) | -98.9 | 13.6 | |

| | | | | |
|---|---|---|---|---|
| #16 | RT change = 1 + TMR * Interference*time + (1+TMR*Interference*time \| subject) | 0 | 0 | Table S3 |
| #17 | VC = 1 + TMR * Interference + time + (1+TMR*Interference \| subject) | -25.7 | 8.6 | |
| #18 | VC = 1 + TMR * Interference*time + (1+TMR*Interference*time \| subject) | 0 | 0 | Table S3 |
| #19 | AC = 1 + TMR * Interference + time + (1+TMR*Interference \| subject) | -1.7 | 5.0 | |
| #20 | AC = 1 + TMR * Interference*time + (1+TMR*Interference*time \| subject) | 0 | 0 | Table S3 |

**Table S3. TMR Effects during Immediate and Delayed Sessions.** This table displays the outcomes of all dependent variables in the main results, considering time as a fixed factor or covariate. When time is treated as a fixed factor, immediate and delayed TMR effects are provided for both interference and non-interference conditions. Bold numbers followed by an asterisk denote significant effects when the highest density interval (HDI) does not overlap with 0. Notably, most TMR effects were observed during the delayed test. The column formula indicates the Bayesian linear mixed model formula that was used, as referenced in Table S2.

<table>
<tr><th rowspan="2">Dependent variables</th><th colspan="2">Interference<br>(cued vs. uncued)</th><th colspan="2">Noninterference<br>(cued vs. uncued)</th><th rowspan="2">Fomular</th></tr>
<tr><th>Immediate</th><th>Delayed</th><th>Immediate</th><th>Delayed</th></tr>
<tr><td rowspan="2">Aversive memory</td><td colspan="2">[-0.82, -0.11]*</td><td colspan="2">[-0.53, 0.64]</td><td>#1</td></tr>
<tr><td>[-0.86, 0.01]</td><td>[-0.87,-0.03]*</td><td>[-0.54, 0.82]</td><td>[-0.61, 0.74]</td><td>#2</td></tr>
<tr><td rowspan="2">Positive memory</td><td colspan="2">[-0.23, 0.65]</td><td colspan="2" rowspan="2"></td><td>#3</td></tr>
<tr><td>[-0.51, 0.79]</td><td>[-0.35, 0.70]</td><td>#4</td></tr>
<tr><td rowspan="2">Aversive intrusions</td><td colspan="2">[-0.40, 0.33]</td><td colspan="2" rowspan="2"></td><td>#5</td></tr>
<tr><td>[-0.64, 0.18]</td><td>[-0.29, 0.68]</td><td>#6</td></tr>
<tr><td rowspan="2">Aversive memory</td><td colspan="2">[-0.68,-0.01]*</td><td colspan="2" rowspan="4"></td><td>#7</td></tr>
<tr><td>[-0.77, 0.09]</td><td>[-0.76, 0.05]</td><td>#8</td></tr>
<tr><td rowspan="2">Positive memory</td><td colspan="2">[-0.28, 0.62]</td><td>#7</td></tr>
<tr><td>[-0.54, 0.95]</td><td>[-0.33, 0.68]</td><td>#8</td></tr>
<tr><td rowspan="2">Aversive memory</td><td colspan="2">[-1.54, -0.24]*</td><td colspan="2" rowspan="4"></td><td>#9</td></tr>
<tr><td>[-2.31, -0.31]*</td><td>[-1.46, 0.14]</td><td>#10</td></tr>
<tr><td rowspan="2">Positive memory (selected)</td><td colspan="2">[0.27, 2.90]</td><td>#9</td></tr>
<tr><td>[-0.02, 3.11]</td><td>[0.31, 3.24]*</td><td>#10</td></tr>
<tr><td rowspan="2">Positive Intrusions</td><td colspan="2">[0.11, 0.65]*</td><td colspan="2">[-0.59, 0.57]</td><td>#11</td></tr>
<tr><td>[-0.09, 0.71]</td><td>[0.10, 0.82]*</td><td>[-0.71, 0.65]</td><td>[-1.12, 0.78]</td><td>#12</td></tr>
<tr><td rowspan="2">Positivity change scores</td><td colspan="2">[0.01, 0.1]</td><td colspan="2">[-0.04, 0.06]</td><td>#13</td></tr>
<tr><td>[0.004, 0.11]*</td><td>[0.0001, 0.11]*</td><td>[-0.05, 0.07]</td><td>[-0.04, 0.07]</td><td>#14</td></tr>
<tr><td rowspan="2">RT changes</td><td colspan="2">[-0.004, 0.04]</td><td colspan="2">[-0.005, 0.04]</td><td>#15</td></tr>
<tr><td>[-0.01, 0.04]</td><td>[-0.01, 0.05]</td><td>[-0.01, 0.05]</td><td>[-0.01, 0.05]</td><td>#16</td></tr>
<tr><td rowspan="2">Valence changes</td><td colspan="2">[-0.31, 0.12]</td><td colspan="2">[-0.30, 0.11]</td><td>#17</td></tr>
<tr><td>[-0.26, 0.25]</td><td>[-0.45, 0.09]</td><td>[-0.26, 0.25]</td><td>[-0.45, 0.09]</td><td>#18</td></tr>
<tr><td rowspan="2">Arousal changes</td><td colspan="2">[-0.31, 0.44]</td><td colspan="2">[-0.30, 0.18]</td><td>#19</td></tr>
<tr><td>[-0.34, 0.43]</td><td>[-0.29, 0.48]</td><td>[-0.46, 0.17]</td><td>[-0.31, 0.32]</td><td>#20</td></tr>
</table>

**Table S4. Time asleep during Day 1 and Day 2, and cueing statistics in each sleep stage during the Day 2 TMR sleep, averaged across participants.** The numbers presented are mean ± SEM. Please note that for some participants, the ground or reference channel became disconnected during the second half of the night due to body movements, resulting in signal loss across all channels. Consequently, these time periods were scored as wake time by YASA. There were five participants on Day 1 and one participant on Day 2 with disconnections; therefore, we only included 30 participants when calculating staging information. However, the cueing number during Day 2 was calculated based on all participants, as there were no disconnections of the ground or reference channel during the first half of the night. A 2 (Days: Day 1 vs. Day 2) by 5 (Sleep Stages: Wake, N1, N2, N3 and REM) repeated-measures ANOVA was conducted on each stages' duration to examine how sleep architecture may differ between Day 1 and Day 2. Results showed that neither the main effect of Days (Day1, Day2) nor the Day by Sleep Stage interaction was significant ($Fs \leqslant 3.74$, $p \geqslant 0.063$).

| **Sleep stage** | **Wake** | **N1** | **N2** | **N3** | **REM** |
|---|---|---|---|---|---|
| Day 1 | 30.0 ± 3.6 | 17.9 ± 1.8 | 228.7 ± 5.8 | 102.7 ± 4.2 | 89.6 ± 3.2 |
| Day 2 | 29.6 ± 3.0 | 19.4 ± 2.2 | 231.9 ± 6.8 | 102.8 ± 4.9 | 104.2 ± 5.2 |
| Cueing number during Day 2 | 0.6 ± 0.2 | 0.0 ± 0.0 | 45.3 ± 12.2 | 460.7 ± 33.3 | 0.9 ± 0.5 |

**Table S5.** The relationship between REM parameters and memory accuracy and memory intrusion in aversive and positive conditions, combining immediate and delayed tests. For aversive and positive memory accuracy, as well as aversive memory intrusion, their baseline performance was controlled using partial Spearman correlation. For positive intrusions, Spearman correlation was employed to examine the relationship between positive intrusions and REM parameters. All p-values were FDR-corrected.

| | **REM parameters** | **Memory accuracy** | ***rho*** | $p_{corrected}$ | **Memory intrusions** | ***rho*** | $p_{corrected}$ |
|---|---|---|---|---|---|---|---|
| Cued | REM percentage | positive | 0.07 | 0.79 | positive | 0.38 | 0.12 |
| | REM Theta power | positive | 0.05 | 0.79 | positive | -0.03 | 0.85 |
| | REM percentage | aversive | 0.18 | 0.79 | aversive | -0.14 | 0.60 |
| | REM Theta power | aversive | -0.05 | 0.79 | aversive | 0.15 | 0.60 |
| Uncued | REM percentage | positive | 0.06 | 0.96 | positive | 0.09 | 0.88 |
| | REM Theta power | positive | -0.03 | 0.96 | positive | -0.03 | 0.88 |
| | REM percentage | aversive | -0.22 | 0.96 | aversive | -0.06 | 0.88 |
| | REM Theta power | aversive | 0.01 | 0.96 | aversive | 0.19 | 0.88 |

**Table S6.** The relationship between REM parameters and positivity change scores, combining immediate and delayed tests. Spearman correlation was used to examine the relationship between positivity change scores and REM parameters. All p-values were FDR-corrected.

| | **REM parameters** | **Positivity change score** | ***rho*** | $\boldsymbol{p}_{\textbf{corrected}}$ |
|---|---|---|---|---|
| Cued | REM percentage | positive | 0.01 | 0.94 |
| | REM Theta power | positive | 0.04 | 0.94 |
| Uncued | REM percentage | positive | 0.04 | 0.81 |
| | REM Theta power | positive | -0.13 | 0.81 |

**Table S7.** The relationship between REM parameters and memory accuracy and memory intrusion in aversive and positive conditions during the immediate test. For aversive and positive memory accuracy, as well as aversive memory intrusion, their baseline performance was controlled using partial Spearman correlation. For positive intrusions, Spearman correlation was employed to examine the relationship between positive intrusions and REM parameters. All p-values were FDR-corrected.

| | **REM parameters** | **Memory accuracy** | ***rho*** | $p_{corrected}$ | **Memory intrusions** | ***rho*** | $p_{corrected}$ |
|---|---|---|---|---|---|---|---|
| Cued | REM percentage | positive | 0.14 | 0.61 | positive | 0.17 | 0.33 |
| | REM Theta power | positive | 0.01 | 0.98 | positive | 0.19 | 0.33 |
| | REM percentage | aversive | 0.20 | 0.61 | aversive | -0.22 | 0.33 |
| | REM Theta power | aversive | -0.13 | 0.61 | aversive | 0.35 | 0.16 |
| Uncued | REM percentage | positive | 0.26 | 0.40 | positive | 0.20 | 0.50 |
| | REM Theta power | positive | 0.04 | 0.81 | positive | 0.05 | 0.93 |
| | REM percentage | aversive | -0.23 | 0.40 | aversive | 0.02 | 0.50 |
| | REM Theta power | aversive | -0.05 | 0.81 | aversive | 0.22 | 0.93 |

**Table S8.** The relationship between REM parameters and memory accuracy and memory intrusion in aversive and positive conditions during the delayed test. For aversive and positive memory accuracy, as well as aversive memory intrusion, their baseline performance was controlled using partial Spearman correlation. For positive intrusions, Spearman correlation was employed to examine the relationship between positive intrusions and REM parameters. All p-values were FDR-corrected.

| | **REM parameters** | **Memory accuracy** | ***rho*** | $p_{corrected}$ | **Memory intrusions** | ***rho*** | $p_{corrected}$ |
|---|---|---|---|---|---|---|---|
| Cued | REM percentage | positive | -0.01 | 0.99 | positive | 0.45 | 0.04 |
| | REM Theta power | positive | 0.09 | 0.99 | positive | -0.23 | 0.48 |
| | REM percentage | aversive | 0.15 | 0.99 | aversive | -0.07 | 0.88 |
| | REM Theta power | aversive | -0.00 | 0.99 | aversive | 0.03 | 0.88 |
| Uncued | REM percentage | positive | -0.15 | 0.76 | positive | -0.03 | 0.99 |
| | REM Theta power | positive | -0.04 | 0.84 | positive | -0.00 | 0.99 |
| | REM percentage | aversive | -0.16 | 0.76 | aversive | -0.13 | 0.99 |
| | REM Theta power | aversive | 0.04 | 0.84 | aversive | 0.06 | 0.99 |

**Table S9.** The relationship between REM parameters and positivity change scores during the immediate and delayed tests. Spearman correlation was used to examine the relationship between positivity change scores and REM parameters. All p-values were FDR-corrected.

| | **REM parameters** | **Positivity change score** | ***rho*** | $p_{corrected}$ |
|---|---|---|---|---|
| Cued immediate | REM percentage | positive | 0.15 | 0.80 |
| | REM Theta power | positive | 0.01 | 0.94 |
| Cued delayed | REM percentage | positive | -0.14 | 0.70 |
| | REM Theta power | positive | 0.07 | 0.70 |
| Uncued immediate | REM percentage | positive | 0.13 | 0.45 |
| | REM Theta power | positive | -0.14 | 0.45 |
| Uncued delayed | REM percentage | positive | -0.05 | 0.91 |
| | REM Theta power | positive | 0.02 | 0.91 |
